# Astrocytic expression of ALS-causative mutant FUS leads to TNFα-dependent neurodegeneration *in vivo*

**DOI:** 10.1101/2021.11.23.469650

**Authors:** Brigid K. Jensen, Kevin J. McAvoy, Nicolette M. Heinsinger, Angelo C. Lepore, Hristelina Ilieva, Aaron R. Haeusler, Davide Trotti, Piera Pasinelli

## Abstract

Genetic mutations that cause Amyotrophic Lateral Sclerosis (ALS), a progressively lethal motor neuron disease, are commonly found in ubiquitously expressed genes. In addition to direct defects within motor neurons, growing evidence suggests that dysfunction of non-neuronal cells is also an important driver of disease. Previously, we demonstrated that mutations in DNA/RNA binding protein Fused in Sarcoma (FUS) induce neurotoxic phenotypes in astrocytes *in vitro,* via activation of the NF-κB pathway and release of pro-inflammatory cytokine TNFα. Here, we developed an intraspinal cord injection model to test whether astrocyte-specific expression of ALS-causative FUS^R521G^ variant (mtFUS) causes neuronal damage *in vivo*. We show that mtFUS expression causes TNFα upregulation, motor function deficits, and spinal motor neuron loss. We further demonstrate a lack of phenotype in TNFα knockout animals expressing mtFUS, and prevention of neurodegeneration in mtFUS-transduced animals through administration of TNFα neutralizing antibodies. Together, these studies strengthen evidence that astrocytes contribute to disease in ALS, establish that FUS-ALS astrocytes induce pathogenic changes to motor neurons *in vivo*, and provide insights identifying FUS-ALS specific potential therapeutic targets.

## INTRODUCTION

Amyotrophic Lateral Sclerosis (ALS) is a progressively fatal neurodegenerative disease caused by degeneration of motor neurons in the brain and spinal cord (Mulder, 1982, Strong et al., 2017). Mutations in the DNA/RNA binding protein Fused in Sarcoma (FUS) account for ∼4% of familial ALS (fALS) and ∼1% of sporadic (sALS) (Renton et al., 2014, Zou et al., 2017). Additionally, mutations in FUS are the most common genetic cause of juvenile ALS (Chen, 2021, Zou et al., 2016). Post-mortem tissues from FUS-ALS patients are marked by cytoplasmic inclusions of FUS in neurons as well as glial cells (Hewitt et al., 2010, Kobayashi et al., 2010, Kwiatkowski et al., 2009, Rademakers et al., 2010, Suzuki et al., 2012). Mislocalization of FUS has also been observed in sALS post-mortem spinal cord tissue and iPSC-derived motor neurons from sALS patients (Fujimori et al., 2018, Ikenaka et al., 2020, Tyzack et al., 2019). Strikingly, altered FUS localization is also found in both cellular and animal models of a distinct genetic form of ALS brought on by mutation of valosin-containing protein (VCP), suggesting that FUS dysregulation may more broadly contribute to disease pathogenesis than is presently recognized (Tyzack et al., 2019). Through DNA, RNA and protein-protein interactions, FUS normally functions in processes of transcription, DNA damage repair, chromosomal stabilization, RNA processing, RNA trafficking and mRNA translation (Deng et al., 2014). The most common pathogenic mutations in FUS, occurring in exons 14 and 15 near the C-terminus of the protein, disrupt nuclear import and lead to cytoplasmic mislocalization, though the degree of nuclear/cytoplasmic localization ranges widely between mutational variants as well as the cell type examined (Dormann et al., 2012, Dormann et al., 2010, Vance et al., 2009, Vance et al., 2013).

ALS pathogenesis is multifactorial, involving repercussions from defects in multiple cell types ultimately converging to damage motor neurons beyond repair (Vucic et al., 2014). While the cell-autonomous effects of FUS mutations on neuronal cells have been analyzed in several studies, considerably less is known about the effect of FUS mutations within non-neuronal cells, and how these changes lead to neuronal damage and death (Machamer et al., 2014, Scekic-Zahirovic et al., 2017, Sharma et al., 2016, Wachter et al., 2015, Xia et al., 2012). Several lines of evidence suggest that astrocytes contribute to neurotoxicity in ALS. Histological studies have reported extensive reactive astrogliosis, gross astrocytic abnormalities, and the presence of protein aggregates, inclusion bodies, and RNA foci in astrocytes within disease-affected areas of patients and animal models of ALS (Averback, 1986, Bruijn et al., 1997, Conlon et al., 2016, Gong et al., 2000, Kamo et al., 1987, Kimura & Budka, 1984, Kushner et al., 1991, Mizielinska et al., 2013, Murayama et al., 1991, Nagy et al., 1994, Popova & Sakharova, 1982). A myriad of molecular changes have also been reported in astrocytes from both ALS patient tissues and animal models (Filipi et al., 2020, Rostalski et al., 2019, Valori et al., 2014). Notably, aberrant transcription factor activity (NF-κB, c-jun, JNK, STAT3), increased inflammatory gene transcripts, and increased cytokine production have been described (Dokalis & Prinz, 2018, Gandelman et al., 2010, Hensley et al., 2003, Migheli et al., 1997, Phatnani et al., 2013, Tong et al., 2013, Tripathi et al., 2017, Tyzack et al., 2017, Wang et al., 2011). Finally, primary astrocytes from transgenic mice carrying ALS-causative gene mutations, primary rodent astrocytes transduced with mutant ALS genes, and astrocytes derived from patient induced pluripotent stem cells have all been found to reduce the viability of wild-type motor neurons (Di Giorgio et al., 2007, Meyer et al., 2014, Nagai et al., 2007, Qian et al., 2017). These studies provide compelling evidence that astrocytes could be a valuable therapeutic target in ALS. However, significant gaps in knowledge currently exist in our understanding of astrocytes as mediators of ALS pathogenesis.

We hypothesized that mutations in FUS could also cause dysregulation of astrocytes, particularly as FUS expression is ubiquitous in the nervous system and FUS participates broadly in essential gene expression regulation processes. Previous studies demonstrate that FUS silencing in rodent astrocytes leads to differential expression of >2000 genes *in vitro* (Fujioka et al., 2013), while CNS silencing of FUS in marmosets induces activation and proliferation of astrocytes absent of any gross neuronal pathology (Endo et al., 2018). Conversely, overexpression of wild-type FUS has been shown to substantially alter the inflammatory profile of mouse and human astrocytes *in vitro* (Ajmone-Cat et al., 2019). Mouse models of FUS also indicate the involvement of non-neuronal cells in disease. Mice harboring the Fus^P525L^ mutation with ubiquitous expression display accelerated disease progression compared with motor-neuron restricted expression (Sharma et al., 2016). In the well-characterized FUS^ΔNLS^ mouse model, excision of mutant FUS specifically from motor neurons was found to delay, but not prevent motor deficits, suggesting that non-cell autonomous effects could be sufficient to drive motor dysfunction (Scekic-Zahirovic et al., 2017). However, it is important to note that the specific contribution of astrocytes to disease course in these models is undetermined. Intriguingly, rare cases of cortical neurodegeneration associated with astrocyte-predominant tauopathy have been linked to a novel FUS variant, however, the pathogenicity of this variant is still uncertain (Ferrer et al., 2015). Taken together, these studies indicate that further examination of the effects of FUS dysregulation in astrocytes is warranted (Scekic-Zahirovic et al., 2017).

Previously, we demonstrated that astrocytes expressing human ALS-linked FUS mutations are toxic to wild-type spinal motor neurons *in vitro* (Kia et al., 2018). We observed that when expressed in astrocytes, mutant FUS induces activation of the transcription factor NF-κB, resulting in elevated production of pro-inflammatory cytokine TNFα (Kia et al., 2018, Qosa et al., 2016). Following binding to receptors on motor neurons, we noted TNFα-induced changes to motor neuron AMPA receptors, which caused sensitization to excitotoxic cell-death (Kia et al., 2018).

To follow up on those studies, here we explored whether mutant FUS expression in astrocytes *in vivo* is also sufficient to induce motor neuron dysfunction and degeneration. We have generated an acute FUS-ALS-astrocyte model in adult mice, using AAV9 to deliver astrocyte-specific promoter (gfa_104_) driven GFP-tagged human FUS. The gfa_104__eGFP, as reference, or gfa_104__eGFP:hFUS^R521G^ (mtFUS) constructs were introduced into the cervical spinal cord of p180 mice by intraspinal injection. The hFUS^R521G^ mutation is a well-established hFUS variant known to cause ALS, localized to the C-terminal PY-NLS (nuclear localization sequence) region of the protein (Dormann & Haass, 2011). It is also one of the variants that we previously utilized in our *in vitro* studies exploring non-cell autonomous effects of astrocyte-expressed mtFUS (Kia et al., 2018, Qosa et al., 2016). We demonstrate that specific expression of hFUS^R521G^ in astrocytes causes upregulation of TNFα, reduction in grip strength and wirehang endurance motor functions, and ultimately a loss of spinal motor neurons. These effects were not seen when TNFα knockout animals underwent the same procedure. We next tested whether FUS-mediated effects could be attenuated through prevention of TNFα signaling in wildtype animals. AAV9 eGFP:hFUS^R521^ intraspinal injections were performed with or without concomitant TNFα-neutralizing antibody (anti-TNFα, α-TNFα). In animals receiving α-TNFα, we observed significant rescue of grip strength performance and preservation of motor neurons. Finally, we designed a protocol which allowed for eGFP:hFUS^R521^ to be expressed for one week prior to administration of TNFα-neutralizing antibody. In these animals, grip strength and motor neurons were preserved at 2-weeks following α-TNFα addition. However, when evaluation of this model was performed at 2-months post-therapeutic administration, the efficacy of this intervention was decreased, with both grip strength performance and motor neuron numbers being significantly reduced.

Overall, these studies further substantiate that astrocytes contribute to disease in genetic models of ALS. We provide novel findings that astrocytic expression of FUS is sufficient to induce pathogenic changes *in vivo* through activation of the NF-κB and TNFα pathway. We also provide evidence that therapeutics designed to diminish activation of this pathway may be effective in reducing motor neuron degeneration in FUS-ALS.

## RESULTS

### Generation and characterization of an acute *in vivo* mouse model for effects of mutant FUS expression in astrocytes

We generated AAV9 encoding either eGFP alone or hFUS^R521G^ fused at the N-terminus to eGFP (eGFP:hFUS^R521G^, hereafter referred to as mtFUS) placed under control of the minimal GFAP promoter element gfa_104_. We performed six injections, bi-laterally, containing 1×10^11^ GC/injection eGFP or mtFUS virus across the C4-C6 region of the spinal cord of p180 mice (**Fig 1A**). We observed high efficiency of AAV9 delivery and enriched targeting to the ventral horn through the intraspinal delivery method (**Fig 1B**). We quantified the transduction efficiency of these vectors by counting eGFP positive cells as a percentage of total cells using nuclear marker DAPI, after 7 days post-injection (DPI) and found 12.7% ± 5.1% of total spinal cord cells transduced (**Fig 1C, Fig EV1A**). In further evaluating only cells within the targeted ventral horn region, 37.8% ± 8.7% transduction efficiency was observed (**Fig 1C**). We also characterized the efficiency of AAV9-GFA104 to transduce astrocytes, by staining spinal cord sections with various cell-type markers at 7 DPI. We analyzed the overlap of eGFP signal with markers for astrocytes (GFAP), neurons (NeuN), microglia (Iba1) and oligodendrocytes (Olig2) (**Fig 1D**). Quantification of this analysis revealed that within the ventral horn, 74.4% ± 6.7% of astrocytes, 6.8% ± 2.3% of neurons, 0% of microglia, and 1.8% ± 1.2% of oligodendrocytes were effectively transduced with the AAV9 viruses (**Fig 1E**). These data demonstrate that intraspinal delivery of AAV9-gfa_104_ robustly transduces astrocytes with minimal leakage into the other cell populations. In evaluating mtFUS localization of transduced cells, we frequently observed a strong nuclear signal along with aggregated cytosolic puncta (**Fig EV1B**). Before analyzing potential mtFUS-mediated effects, to better characterize this model, we assessed expression of eGFP and mtFUS over time. No differences in overall body weight were observed following a 4-week observation period post-surgery (**Fig S1C**). Robust eGFP expression was observed in as little as 3 days in both GFP and mtFUS animals when evaluated by immunofluorescent staining (**Fig EV2A**). At 14 days post-injection, GFP levels were still robustly and significantly expressed in both GFP and mtFUS cohort groups when evaluated by immunofluorescent staining of spinal cords, qRT-PCR mRNA analysis, and western blotting for GFP protein (**Fig EV2A-C**). Evaluation of FUS levels in these mice revealed similar levels of total FUS protein when evaluated by immunofluorescence and western blotting (**Fig EV2D-F**). Of note, in qRT-PCR analysis, only mtFUS animals showed mRNA specific to the human FUS sequence (*p* < .01) (**Fig EV2E**). Similarly, only mtFUS animals had a second FUS band on the western blot corresponding to the GFP-FUS fusion protein (**Fig EV2F**). As a result of these assessments, we concluded that this model was appropriate for studying the potential effects of astrocytic mtFUS expression *in vivo*.

**Figure 1:**
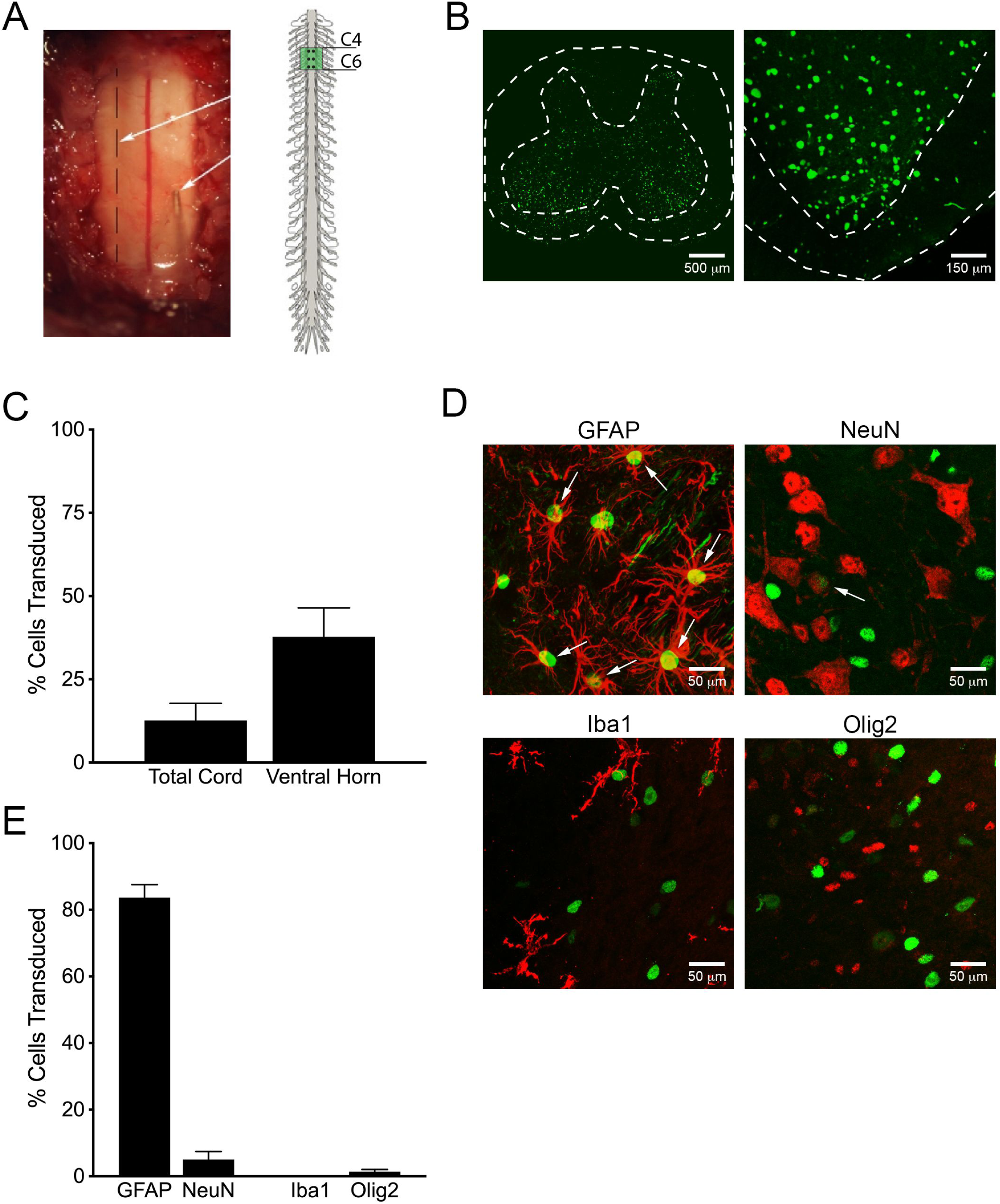
Generation and characterization of an acute *in vivo* mouse model for effects of mutant FUS expression in astrocytes. A) Model design: Left: Image of mouse cervical spinal region indicating the AAV delivery site. Middle: Cartoon of spinal cord region within the context of the total cord. Six injections are delivered bi-laterally from C4 to C6 region. B) Representative transduction profile showing 10x magnification image (scale bar indicates 500 μm) and cropped 20x magnification image (scale bar indicates 150 μm) of eGFP:hFUS^R521G^ (mtFUS) in a 30 μm spinal cord slice within the transduction region after six 1×10^^11^ GC injections in p180 female wildtype mice. C) Transduction efficiency of AAV9-gfa104 after 1 week of transduction. Quantification of GFP^+^ cells within DAPI masked region of spinal cord, showing percentages of total cord, and ventral horn cells transduced. Quantification performed using image calculator functions in ImageJ revealed approximately 12.7% of total cells, and 37.8% of ventral horn cells were effectively transduced with virus using these dosing parameters. Data presented as mean ± SEM. D) Representative immunohistochemical staining of cell-type specific markers in 30 μm spinal cord sections of mice 1-week post-injection, 60x magnification, scale bar indicates 50 μm. The cell types tested are astrocytes (GFAP), neurons (NeuN), microglia (Iba1) and oligodendrocytes (Olig2). Cell type specific markers are shown in red, with GFP^+^ cells indicating those which were transduced with virus. E) Quantification of the percentage GFP^+^ cells out of total cells labelled with each cell marker. 74.4% of astrocytes, 6.8% of neurons, 0% of microglial, and 1.8% of oligodendrocytes were effectively transduced using these dosing parameters. Data presented as mean ± SEM. For these evaluations n = 3 mice per condition (eGFP or mtFUS) were assessed, with 5-10 spinal cord sections analyzed per each subject.

### Motor deficits and loss of motor neurons occur when mtFUS is expressed in astrocytes

An advantage of utilizing intraspinal injection sites spanning the C4-C6 region of the mouse spinal cord is that motor neuron loss in this region has the potential to impact readily measurable forelimb motor behaviors (Bacskai et al., 2013). Assessment of these behaviors enabled a first-pass evaluation of whether mtFUS expression in astrocytes *in vivo* produced motor deficits over time. At 14 days post-injection, we noticed overt weakness of the mouse forelimb in mtFUS expressing animals, but not eGFP alone. We measured a permanent reduction in both grip strength and wire hang endurance, beginning at 2 weeks, in mtFUS animals but not eGFP-alone (**Fig. 2A-B**). At 2 weeks, mtFUS animals showed a wire hang latency of 30.83 seconds ± 6.9 seconds, compared with eGFP animals who had a wire hang latency of 81.7 seconds ± 13.4 seconds (*p* < .01). Similarly, at 2 months this phenotype was maintained, with mtFUS animals having a wire hang endurance of 23.4 seconds ± 7.8 seconds, compared with eGFP animals with a wire hang latency of 91.7 seconds ± 8.9 seconds (*p* < .0001) (**Fig. 2A**). As a second measure of motor performance, grip strength was also measured at these timepoints. At 2 weeks mtFUS animals had a grip strength of 0.10 kg compared with eGFP animals with a grip strength of 0.16 kg (*p* < .01), and at 2 months mtFUS animals had a grip strength of 0.08 kg compared with the 0.18 kg grip strength of eGFP controls (*p* < .001) (**Fig. 2B**). We next sought to determine whether this loss of motor function correlated with a loss of motor neurons. We counted motor neurons in spinal cord sections from these mice using two motor neuron markers: matrix metalloproteinase 9 (MMP9), which is a specific marker for the large motor neuron pool most vulnerable in ALS, and choline acetyltransferase (ChAT) which marks all cholinergic motor neurons. At the 3-day post-injection timepoint prior to overt symptoms, we do not observe a loss of MMP9^+^ cells **(Fig EV3A).** However, at the 2-week timepoint when we do observe motor symptoms, there is a loss of MMP9^+^ cells in mtFUS mice, corresponding to only 59.6% ± 8.5% of that found in sham surgery (*p* < .01) and eGFP animals (*p* < .05) (**Fig 2C, E**). Further, we see a similar temporal pattern of ChAT^+^ cell loss in mtFUS expressing animals, with similar ChAT levels among groups at 3 days post-injection, which is reduced significantly in mtFUS mice at the 2-week time point, falling to 51.5% ± 3.0% of sham surgery (*p* < .05) and eGFP animals (*p* < .05) **(Fig 2D,F, Fig EV3B)**. These results provide evidence that astrocyte-restricted mtFUS expression in adult mice can cause motor neuron dysfunction and death in as little as 2 weeks.

**Figure 2.**
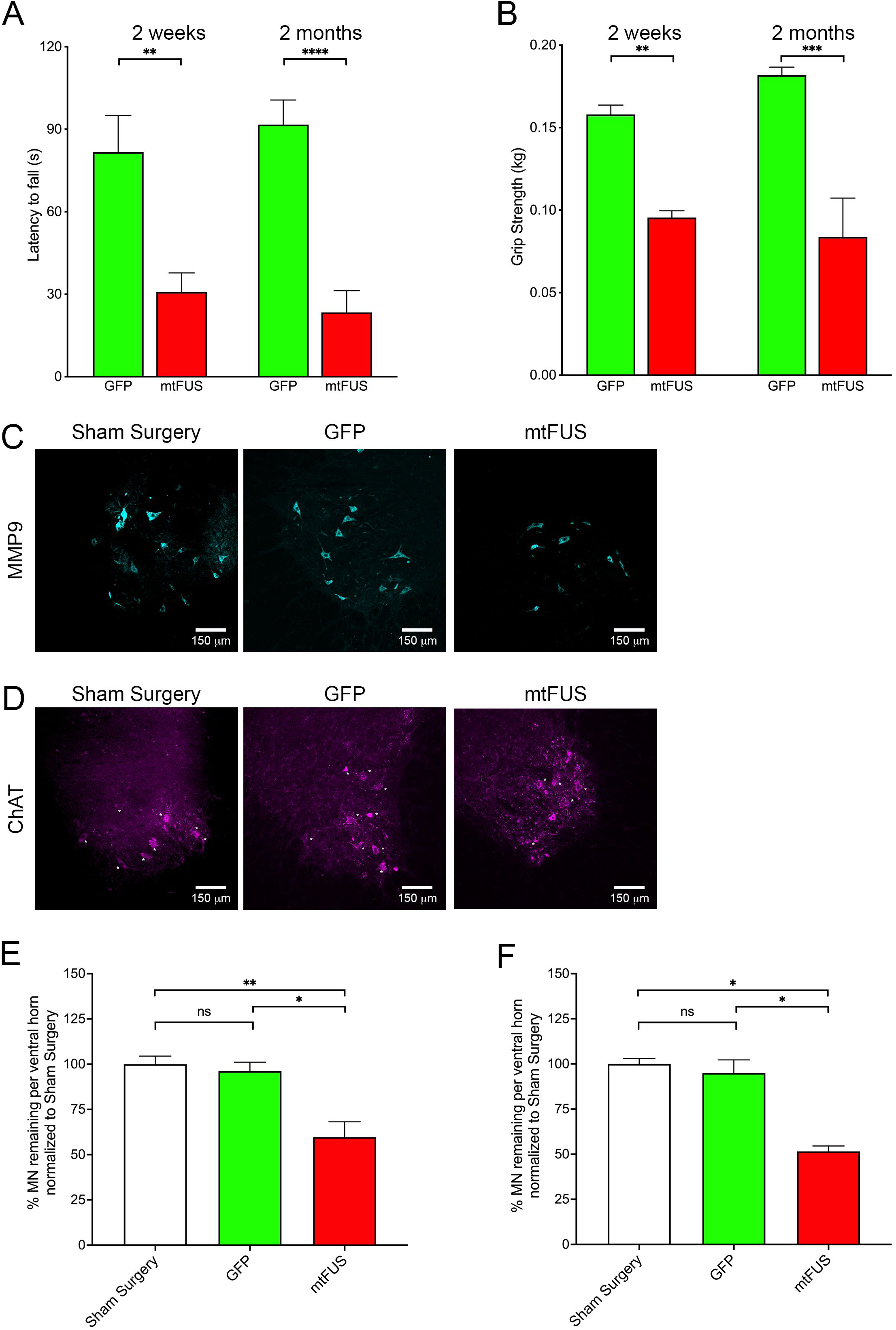
Motor deficits and loss of motor neurons occur when mtFUS is expressed in astrocytes. A) Quantification of motor behavior by wire hang latency, in eGFP and mtFUS mice at 2-weeks and 2-months months post-injection. At 2-weeks, animals expressing mtFUS had a significantly reduced wire hang latency, compared with eGFP animals. This was a permanent effect, as at 2-months, mtFUS animals again showed a significantly reduced latency compared with eGFP animals. Data presented as mean ± SEM, n = 5 animals per timepoint per condition. Dunnett’s multiple comparison test determined statistical significance for comparison of GFP versus mtFUS at each timepoint, ** *p* < .01, **** *p* < .0001. B) Forelimb grip-strength was also assessed in eGFP and mtFUS animals at 2 weeks post-injection and 2-months post injection. At 2-weeks, mtFUS expressing animals had a significantly reduced grip strength compared with eGFP animals. This also was a permanent effect, at 2-months, mtFUS animals again showed a significantly reduced grip strength compared with eGFP animals. Data presented as mean ± SEM, n = 5 animals per timepoint per condition. Dunnett’s multiple comparison test determined statistical significance for comparison of GFP versus mtFUS at each timepoint, ** *p* < .01, *** *p* < .001. C) Immunohistochemical staining for MMP9, a marker for vulnerable large motor-neurons in ALS, in spinal cord sections at 14 days post-injection. Sham surgery and eGFP expressing animals show similar numbers of MMP9^+^ neurons visualized in cyan, while in mtFUS animals there are fewer positive cells. Representative fields from 60x magnification z-stack confocal images, scale bar indicates 150 μm. D) Immunohistochemical staining for the motor neuron marker ChAT at 14 days post-injection. Sham surgery and eGFP expressing animals show similar numbers of ChAT^+^ neurons visualized in magenta and indicated by *, while in mtFUS animals there are fewer positive cells. Representative fields from 60x magnification z-stack confocal images, scale bar indicates 150 μm. E) MMP9^+^ motor neurons were quantified within the ventral horn of spinal cord sections by manual counting of immunopositive cells within the ventral horn region. Analysis revealed a significant reduction in MMP9^+^ cells in mtFUS expressing animals compared with either sham surgery controls (\*\**p* < .01) or eGFP expressing animals (\**p* < .05). Data presented as mean ± SEM, n = 4 animals per condition, m= 4 ventral horns per animal. Sidak’s multiple comparison test determined statistical significance. F) ChAT^+^ motor neurons were also quantified within the ventral horn of spinal cord sections by manual counting of immunopositive cells within the ventral horn region. Quantification showed a significant reduction in ChAT^+^ cells in mtFUS expressing animals compared with either sham surgery controls (\**p* < .05) or eGFP expressing animals (\**p* < .05). Data presented as mean ± SEM, n = 4 animals per condition, m= 4 ventral horns per animal. Sidak’s multiple comparison test determined statistical significance.

### TNFα levels are elevated in mtFUS mice

Next, we sought to determine whether TNFα signaling was altered in mtFUS animals as we had previously observed *in vitro (Kia et al., 2018).* Using qRT-PCR analysis on cervical spinal cord sections of animals 14 DPI, we found significantly elevated TNFα mRNA expression in mtFUS mice, with levels 2.6 ± 0.41-fold higher than animals expressing eGFP (*p* < .01) (**Fig 3A**). Additionally, the soluble fraction from cervical spinal cord homogenates was analyzed with a highly sensitive TNFα ELISA, which revealed a significantly increased level of secreted TNFα protein in mtFUS animals compared with eGFP control animals, with 3.5 ± 0.14 pg/mL detected in eGFP animals, and 5.3 ± 0.7 pg/mL detected in mtFUS animals (*p* < .05) (**Fig 3B**). This elevation of TNFα corresponded with increased astrocyte reactivity as measured by glial fibrillary acidic protein (GFAP) at both the mRNA level where a 3.0 ± 0.46-fold increase was seen (*p* < .001) (**Fig EV4A**) and at the protein level through enhanced signal via immunofluorescent staining when compared with eGFP control animals (**Fig EV4B**). These data suggest that astrocytic mtFUS expression leads to pro-inflammatory changes *in vivo*, which include elevated TNFα levels and astrocytic reactivity in the surrounding region.

**Figure 3.**
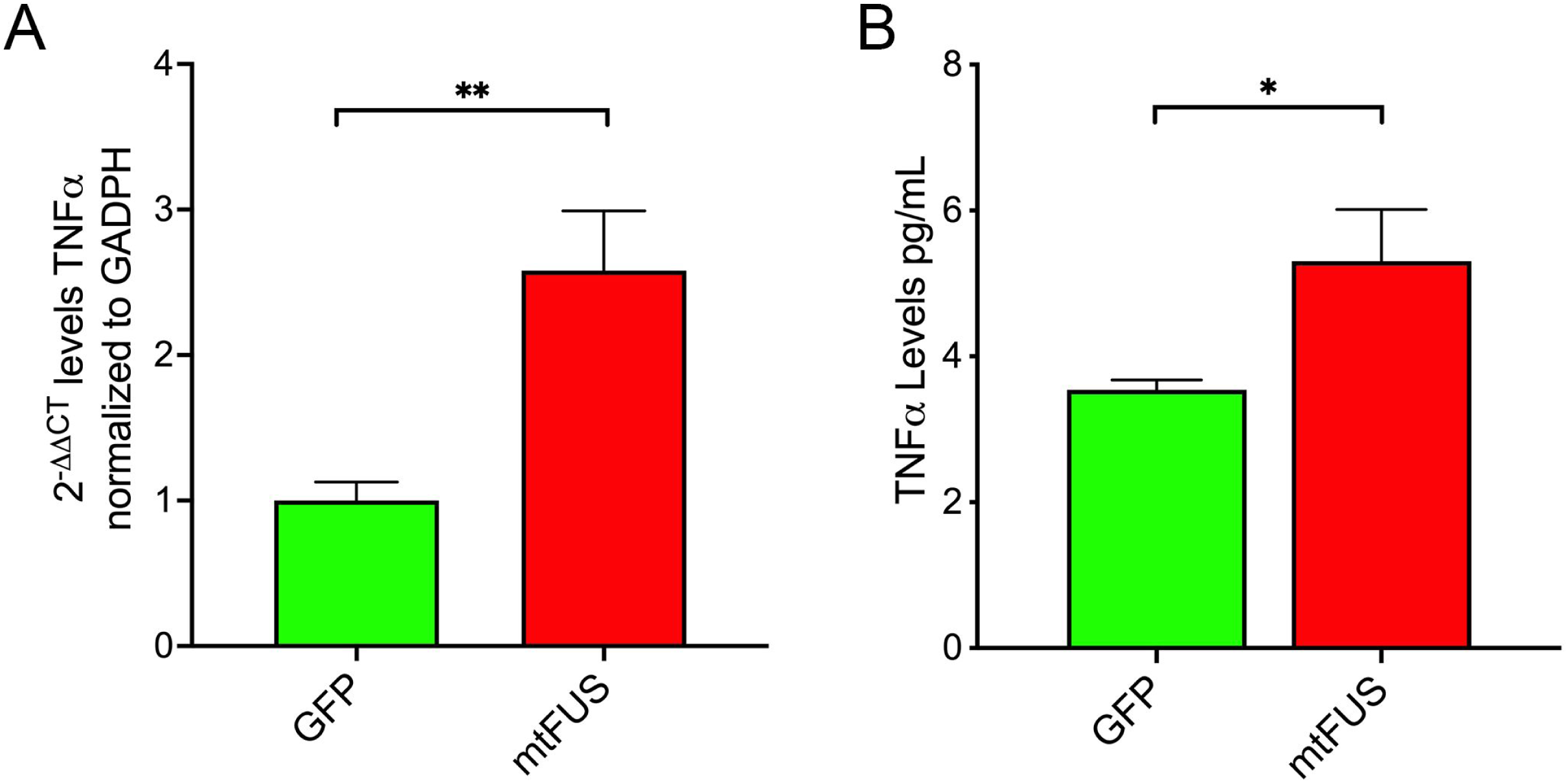
TNFα levels are elevated in mtFUS mice. A) qPCR analysis of TNFα levels at 14 days post-injection indicates a significantly elevated level of expression in mtFUS expressing animals compared with eGFP controls. Data shown as relative fold-changes compared to GFP alone, and are presented as mean ± SEM, Student’s t test determined statistical significance, n=9 animals per group, \*\*\**p* < 0.01. B) A highly sensitive quantitative TNFα ELISA was performed to assess soluble TNFα levels in animals expressing GFP or mtFUS for 2-weeks. A significant increase in TNFα was detected in mtFUS animals. Data presented as mean ± SEM, Student’s *t*-test determined statistical significance, n=6 animals per group, * *p* < 0.05.

### TNFα knockout animals do not display motor deficits or loss of motor neurons when mtFUS is expressed in astrocytes

We next sought to explore whether the behavioral and motor neuron phenotypes that we observed were explicitly caused by the action of secreted TNFα *in vivo*. To study this, we repeated intraspinal delivery of eGFP or mtFUS in p180 homozygous TNFα knockout (TNF KO) and paired wildtype (TNF ^+/+^) animals. At 2 weeks post-injection, these animals were evaluated for grip strength and assessed for spinal cord motor neuron number by MMP9 and ChAT staining as done in Figure 2. In striking contrast to the paired wildtype (TNF^+/+^) controls expressing mtFUS, which displayed a significant reduction from 0.22 kg to 0.13 kg grip strength (*p* < .01), TNFα knockout animals receiving mtFUS did not have any deficits in grip strength, with comparable measurements of 0.22kg in both sham surgery and mtFUS expressing groups (**Fig 4A**). Moreover, while wildtype TNF^+/+^-mtFUS animals again showed reductions in MMP9^+^ and ChAT^+^ cells (to 52.4% and 57.7% in mtFUS versus sham surgery wildtype animals respectively (*p* < .01)), we also did not observe any decrease in MMP9^+^ or ChAT^+^ motor neurons in the spinal cord of TNFα knockout animals, with values of 96.9% MMP9^+^ and 97.0% ChAT^+^ cells respectively in mtFUS compared with sham surgery animals (**Fig 4B-E**). Genotype was confirmed in these animals by PCR analysis to verify TNFα knockout (**Fig EV5A**). Comparable levels of mtFUS expression was also validated in wildtype TNF^+/+^ and TNFα knockout animals receiving mtFUS virus by qRT-PCR analysis (*p* < .001) (**Fig EV5B**). Overall, these findings suggest that TNFα signaling is substantially responsible for the observed motor behavioral deficits and motor neuron cell death that is caused in our mouse model.

**Figure 4:**
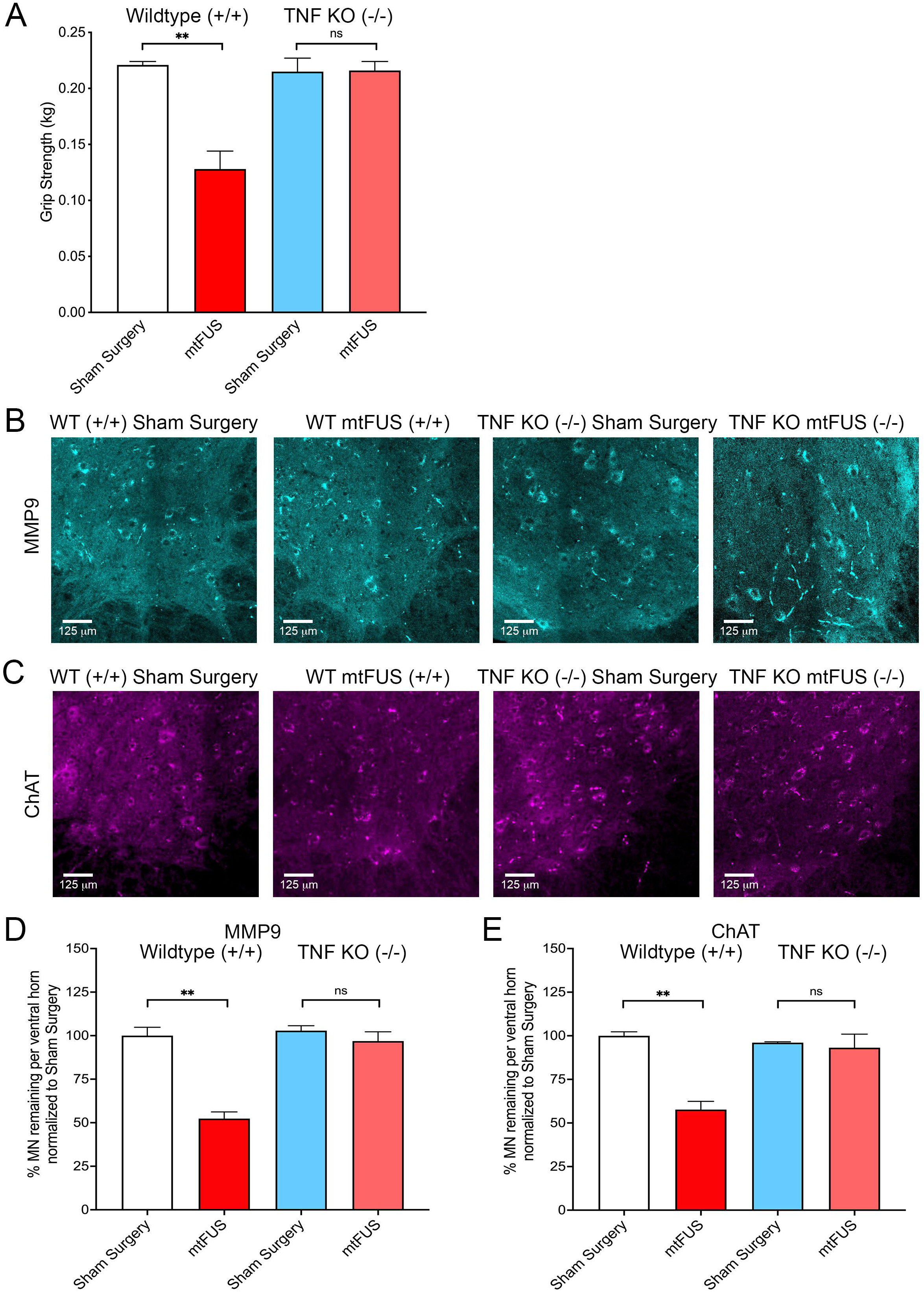
TNFα knockout animals do not display motor deficits or loss of motor neurons when mtFUS is expressed in astrocytes. A) Forelimb grip-strength was quantified in an age-matched paired cohort of wildtype and TNFα knockout (TNF KO) mice at 2-weeks following either sham surgery or mtFUS expression. At this time, mtFUS expressing animals in the wildtype (TNF^+/+^) cohort had a significantly reduced grip strength of 0.13 kg, compared with sham surgery animals with a grip strength of 0.22kg. In contrast, in the TNF KO cohort, animals displayed comparable grip strength values of 0.22kg regardless of whether they had undergone sham surgery or were expressing mtFUS. Data presented as mean ± SEM, n = 3 animals per timepoint per condition. Dunnett’s multiple comparison test determined statistical significance for comparison of sham versus mtFUS for each genotype, ** *p* < .01. B) Representative immunohistochemical staining for MMP9 (cyan) in spinal cord sections from wildtype and TNF KO animals having undergone sham surgery or expressing mtFUS at 14 days post-injection. In wildtype animals, mtFUS expressing animals show a reduction in numbers of MMP9^+^ neurons. In TNF KO animals, comparable numbers of MMP9^+^ cells are seen both in the sham surgery and mtFUS conditions. Representative fields from 60x magnification z-stack confocal images, scale bar indicates 125 μm. C) Representative immunohistochemical staining for ChAT (magenta) at 14 days post-injection from wildtype and TNF KO animals having undergone sham surgery or expressing mtFUS. In wildtype animals, mtFUS expressing animals show a reduction in numbers of ChAT^+^ neurons. In TNF KO animals, comparable numbers of ChAT^+^ cells are seen both in the sham surgery and mtFUS conditions. Representative fields from 60x magnification z-stack confocal images, scale bar indicates 125 μm. D) Quantification of MMP9^+^ motor neurons was performed as in Figure 2. Analysis revealed a significant reduction in MMP9^+^ cells in wildtype animals expressing mtFUS compared to wildtype sham surgery animals (** *p* < .01). In contrast, comparable numbers of MMP9^+^ cells were found in both sham surgery and mtFUS animals in the TNF KO cohort. Data presented as mean ± SEM, n = 3 animals per condition, m= 4 ventral horns per animal. Sidak’s multiple comparison test determined statistical significance. E) Quantification of ChAT^+^ motor neurons was performed as in Figure 2. A significant decrease in ChAT^+^ motor neurons was observed in wildtype animals expressing mtFUS compared to their paired sham surgical controls (** *p* < .01). However, similar numbers of ChAT^+^ cells were found in both the sham surgery and mtFUS groups for the TNF KO cohort. Data presented as mean ± SEM, n = 3 animals per condition, m= 4 ventral horns per animal. Sidak’s multiple comparison test determined statistical significance.

### α−TNFα neutralizing antibody application at the time of mtFUS injection prevents motor neuron death and functional deficits in mtFUS expressing animals

Next, we tested whether pharmacological manipulation of TNFα could also prevent motor dysfunction and motor neuron death in our mouse model. For this study, we have applied a TNFα neutralizing antibody at an effective dosing of 2 mg/kg, as has been reported in other *in vivo* experimental models (Finsterbusch et al., 2016, Via et al., 2001). First, we evaluated the consequences of co-administering the TNFα neutralizing antibody at the time of viral injection. In this paradigm, antibody was co-injected with AAV, while maintaining the same 1μL total injection volume and viral titer as in previous cohorts. At two weeks post-injection, animals receiving mtFUS virus + TNFα neutralizing antibody (α-TNFα) displayed similar grip strength to their control sham surgery counterparts (0.14 kg compared to 0.16 kg), while mtFUS only animals demonstrated the same significant grip strength deficit as observed in earlier cohorts (0.11 kg) (*p* < .01) (**Fig 5A**). Additionally, animals which received mtFUS+ α-TNFα had preserved motor neuron numbers comparable to sham surgical controls (MMP9^+^ 85.3% and ChAT 100.2% respectively), while animals which received mtFUS alone showed significant reductions as previously observed (MMP9^+^ 61.7%, sham vs mtFUS *p* < .0001, mtFUS+ α-TNFα vs mtFUS *p* < .01; ChAT^+^ 71.3%, sham vs mtFUS *p* < .01, mtFUS+ α-TNFα vs mtFUS *p* < .01)) (**Fig 5B-C**). In this surgical paradigm, we have confirmed that α-TNFα had no effect on levels of mtFUS expression by qRT-PCR analysis (*p* < .01) (**Fig EV6A**). From this cohort of animals, we have observed that when α-TNFα is administered concomitant with intraspinal mtFUS virus delivery, effects on motor neuron dysfunction and death are greatly attenuated, such that animals display no overt motor deficits or loss of motor neurons.

**Figure 5:**
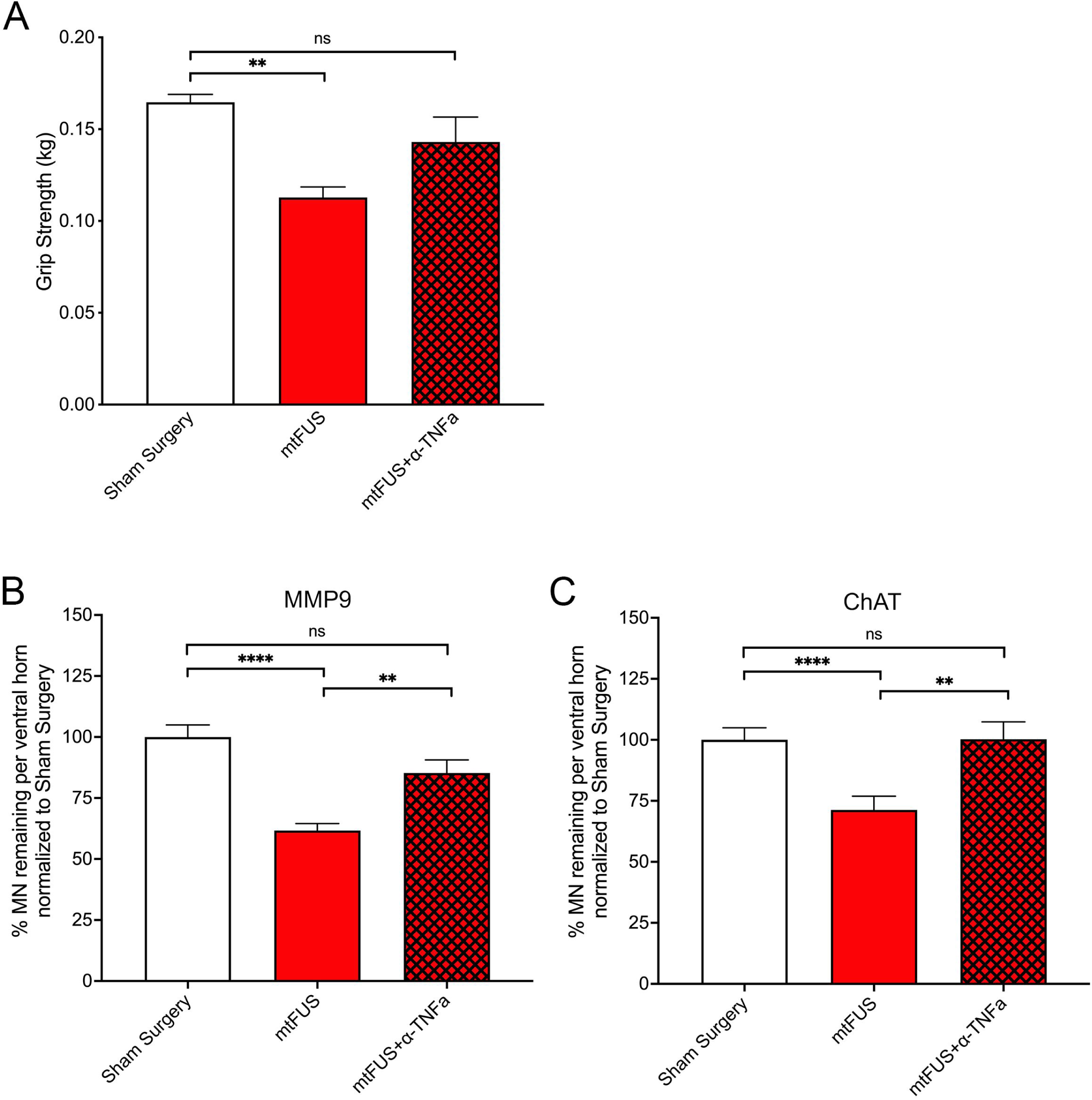
α−TNFα neutralizing antibody application at the time of mtFUS injection prevents motor neuron death and functional deficits in mtFUS expressing animals. A) Forelimb grip-strength was quantified in a cohort containing sham surgery animals, mtFUS expressing animals, and mtFUS expressing animals that received a-TNFa neutralizing antibody (mtFUS+ α-TNFα) at the time of viral delivery, two weeks after intraspinal injections. mtFUS expressing animals had a significantly reduced grip strength compared with sham surgery animals, as had been previously seen. In the mtFUS+ α-TNFα cohort however, animals displayed grip strength values which did not statistically differ compared with the sham surgery group. Data presented as mean ± SEM, n = 5 animals per timepoint per condition. One-way ANOVA with Sidak’s multiple comparison test determined statistical significance, ** *p* < .01. B) Quantification of MMP9^+^ motor neurons was performed as in Figure 2. As previously noted, analysis revealed a significant reduction in MMP9^+^ cells in wildtype animals expressing mtFUS compared to wildtype sham surgery animals (**** *p* < .0001). In contrast, mtFUS expressing animals which received α-TNFα neutralizing antibody showed a significant restoration of MMP9^+^ compared with the mtFUS only group (** *p* < .01) and had levels that were not statistically altered from the sham surgery cohort. Data presented as mean ± SEM, n = 6 animals per condition, m= 4 ventral horns per animal. One-way ANOA with Sidak’s multiple comparison test determined statistical significance. C) Quantification of ChAT^+^ motor neurons was performed as in Figure 2. A significant decrease in ChAT^+^ motor neurons was again seen in animals expressing mtFUS compared to the sham surgical controls (** *p* < .01). mtFUS+ α-TNFα animals showed a significant restoration of ChAT^+^ compared with the mtFUS only group (** *p* < .01) and had levels that were not statistically altered from the sham surgery cohort. Data presented as mean ± SEM, n = 6 animals per condition, m= 4 ventral horns per animal. One-way ANOVA with Sidak’s multiple comparison test determined statistical significance.

### α−TNFα neutralizing antibody application one week following mtFUS injection prevents motor neuron death and functional deficits in mtFUS expressing animals for a period of time

Finally, we assessed the effects of administering TNFα neutralizing antibody after a period of mtFUS expression, to better understand if this could be a viable therapeutic approach when mtFUS is already present and evoking cellular effects. In designing this paradigm, we chose to investigate outcomes of a single application of neutralizing antibody to quickly ascertain whether blocking TNFα-mediated effects would have any utility if mtFUS was already perturbing the system. In this surgical paradigm, mtFUS intraspinal injections were performed as in all previous cohorts, with a non-adhesive dressing applied over the site of spinal cord injections just prior to surgical closure. Following one week of recovery and mtFUS expression, animals underwent a second surgery in which the spinal cord was again exposed at the same location, the dressing was removed, and was replaced with an absorbable gelatin sponge soaked in sterile saline +/- α-TNFα. Animals were allowed to recover and were assessed for grip strength and motor neuron numbers at a 2-week endpoint following this second surgery. At 2 weeks following α-TNFα administration we observed grip strength, MMP9^+^, and ChAT^+^ neurons which were comparable to sham surgery animals and which were significantly restored compared to mtFUS animals which had not received neutralizing antibody (**Fig 6**). At this timepoint, mtFUS only expressing animals displayed a significant reduction of grip strength from 0.18 kg to 0.10 kg (*p* < .0001), however, animals receiving α-TNFα following one week of mtFUS expression had a significantly elevated measured grip strength of 0.17 kg (*p* < .01) (**Fig 6A**). Similarly, while MMP9^+^ cells fell to 45.8% in mtFUS only animals at 2 weeks (p < .001), those also receiving α-TNFα still had 91.9% of cells compared with sham surgery animals, a significant increase over the mtFUS animals (*p* < .01) (**Fig 6C**). Likewise, when ChAT^+^ cells were evaluated, mtFUS only animals had a significant reduction in cell number (47.8%) (*p* < .001), mtFUS + α-TNFα animals showed a significant recovery of cell number, comparable to that of sham surgery animals (94.2%, *p* <.01) (**Fig 6E**). Given this striking result following 2-weeks of α-TNFα application, we extended our trials in an additional cohort to determine if animals retained motor neuron functionality and viability 2-months after delivery of the single-dose application of α-TNFα neutralizing antibody. In contrast to our 2-week results, at 2 months grip strength was reduced from 0.16 kg to 0.11 kg and no longer had a significant recovery from the mtFUS animals (0.08 kg) (**Fig 6B**). Additionally, both MMP9^+^ (66.8%) and ChAT^+^ cells (56.7%) were significantly reduced compared to sham surgery (*p* < .0001), and instead were comparable to mtFUS animals that had not received neutralizing antibody (**Fig 6D,F**). In this surgical paradigm, levels of mtFUS expression were again confirmed to be comparable in mtFUS and mtFUS+ α-TNFα groups by qRT-PCR analysis at both 2-week and 2-month evaluation timepoints (**Fig EV6B**). In conclusion, we have found that a single dosing of α-TNFα prevented astrocytic mtFUS-mediated deficits in motor neuron function and survival for a period of time extending to two weeks post α-TNFα administration. While we anticipate that loss of efficacy of the α-TNFα antibody by our 2-month evaluation timepoint is due to usage and/or diffusion to levels that are sub-threshold to prevent TNFα signaling, we cannot evaluate this directly using our present model. We are able to conclude however, that once the efficacy of α-TNFα treatment is lost the neurodegenerative process continues to proceed, indicating the mtFUS expression in astrocytes is indeed driving these deleterious consequences.

**Figure 6:**
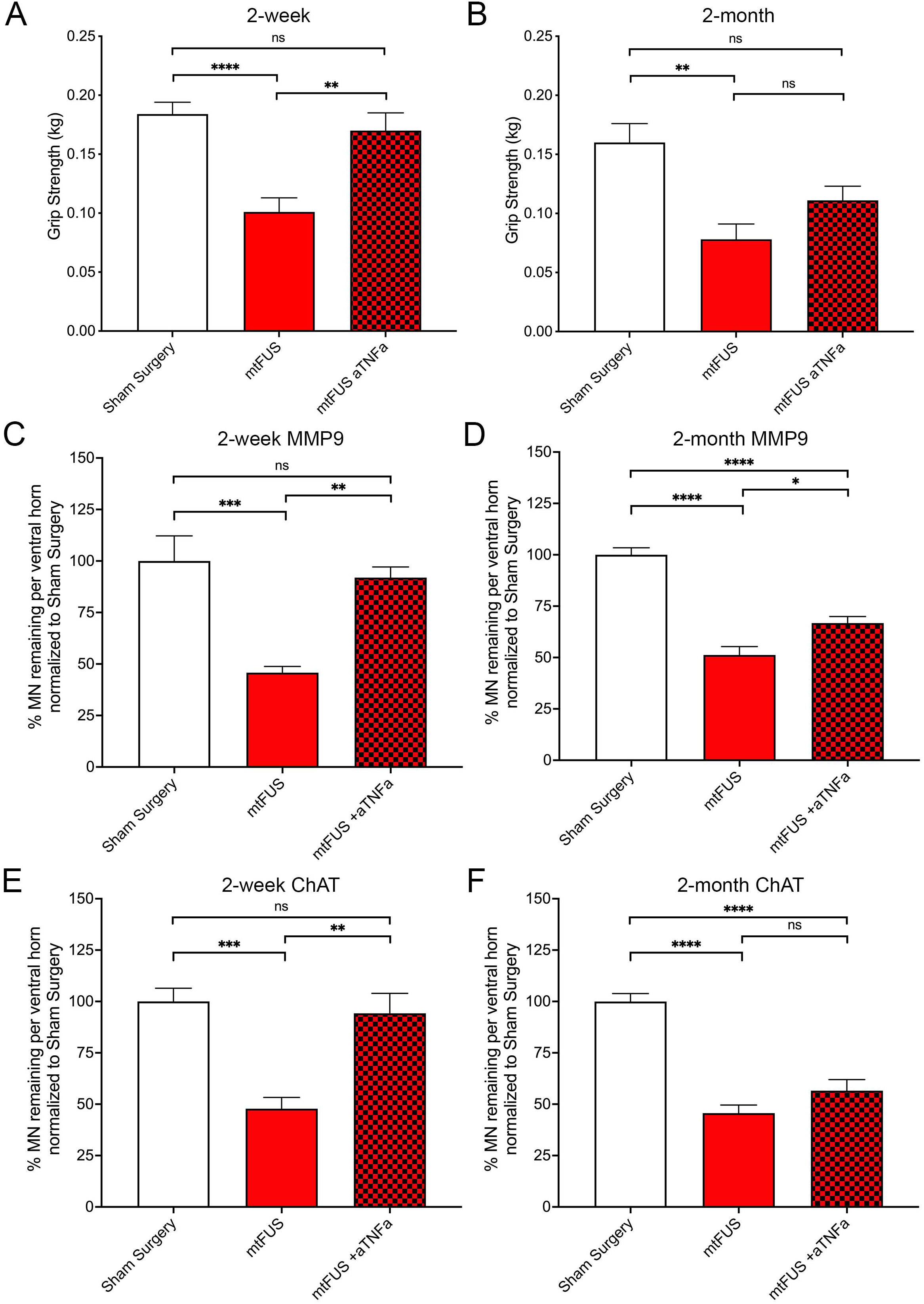
α−TNFα neutralizing antibody application one week following mtFUS injection prevents motor neuron death and functional deficits in mtFUS expressing animals for a period of time. A) In the paradigm of viral infection with mtFUS, followed a week later by α-TNFα neutralizing antibody application, forelimb grip-strength was quantified two weeks after administration of α-TNFα. mtFUS expressing animals had a significantly reduced grip strength compared with sham surgery animals (**** *p* < .0001). In the mtFUS+ α-TNFα cohort, animals displayed grip strength which significantly elevated compared with the mtFUS only animals (** *p* < .01), and which did not statistically differ compared with the sham surgery group. Data presented as mean ± SEM, n = 6 animals per timepoint per condition. One-way ANOVA with Sidak’s multiple comparison test determined statistical significance. B) In this same paradigm, forelimb grip-strength was quantified two months after application of α-TNFα. mtFUS expressing animals had a significantly reduced grip strength of 0.08 kg compared with sham surgery animals with a grip strength of 0.16kg (** *p* < .01). By this time, mtFUS+ α-TNFα cohort also displayed grip strength reduction, with values of 0.11kg, which trended towards a significant reduction compared with sham surgery controls (*p* = 0.13), and which not statistically different compared with the mtFUS only animals. Data presented as mean ± SEM, n = 6 animals per timepoint per condition. One-way ANOVA with Sidak’s multiple comparison test determined statistical significance. C) Quantification of MMP9^+^ motor neurons was performed as in Figure 2 for our delayed α-TNFα surgical group. At two weeks following neutralizing antibody application, analysis revealed a significant reduction in MMP9^+^ cells in animals expressing mtFUS only compared to sham surgery animals (*** *p* < .001). In contrast, mtFUS expressing animals which received α-TNFα neutralizing antibody showed a significant restoration of MMP9^+^ compared with the mtFUS only group (** *p* < .01) and had levels that were not statistically altered from the sham surgery cohort. Data presented as mean ± SEM, n = 6 animals per condition, m= 4 ventral horns per animal. One-way ANOVA with Sidak’s multiple comparison test determined statistical significance. D) Quantification of MMP9^+^ motor neurons was performed as in Figure 2 for our delayed α-TNFα surgical group at two months following α-TNFα antibody application. Analysis showed a significant reduction in MMP9^+^ cells in animals expressing mtFUS only compared to sham surgery animals (**** *p* < .0001). By this time, mtFUS expressing animals which received α-TNFα neutralizing antibody also showed a significant reduction in MMP9^+^ cells compared with the sham surgery group (**** *p* < .0001) and had levels that were only slightly elevated compared with the mtFUS only group (* *p* < .05). Data presented as mean ± SEM, n = 6 animals per condition, m= 4 ventral horns per animal. One-way ANOVA with Sidak’s multiple comparison test determined statistical significance. E) Quantification of ChAT^+^ motor neurons was performed as in Figure 2 for our delayed α-TNFα surgical group. At two weeks following neutralizing antibody application, analysis revealed a significant reduction in ChAT^+^ cells in animals expressing mtFUS only compared to sham surgery animals (*** *p* < .001). In contrast, mtFUS expressing animals which received α-TNFα neutralizing antibody showed a significant restoration of ChAT^+^ compared with the mtFUS only group (** *p* < .01) and had levels that were not statistically altered from the sham surgery cohort. Data presented as mean ± SEM, n = 6 animals per condition, m= 4 ventral horns per animal. One-way ANOA with Sidak’s multiple comparison test determined statistical significance. F) Quantification of ChAT^+^ motor neurons was performed as in Figure 2 for our delayed α-TNFα surgical group at two months following α-TNFα antibody application. Analysis showed a significant reduction in ChAT^+^ cells in animals expressing mtFUS only compared to sham surgery animals (**** *p* < .0001). By this time, mtFUS expressing animals which received α-TNFα neutralizing antibody also showed a significant reduction in ChAT^+^ cells compared with the sham surgery group (**** *p* < .0001) and had levels that were comparable to the mtFUS only group. Data presented as mean ± SEM, n = 6 animals per condition, m= 4 ventral horns per animal. One-way ANOA with Sidak’s multiple comparison test determined statistical significance.

## DISCUSSION

Expanding upon our previous *in vitro* work, (57, 58), in this study we demonstrate that expression of mtFUS in astrocytes alone can drive motor neuron dysfunction and death *in vivo*. Here, we have developed an acute *in vivo* astrocytic expression model in adult mice, using intraspinal AAV injections targeted to spinal levels associated with forelimb motor function. Through this method, we show that astrocytes expressing mtFUS become neurotoxic and induce focal motor neuron dysfunction and death, impairing motor behavior. To our knowledge, this is the first *in vivo* reporting of such astrocytic non-cell autonomous processes in a FUS-ALS model. We also provide evidence that that these effects occur through a TNFα-mediated mechanism, identifying an interesting, and potentially mtFUS-specific, toxic pathway to target therapeutically. Indeed, through injection of an α-TNFα neutralizing antibody we show that it is possible to prevent or block motor neuron degeneration and associated behavioral deficits in mtFUS-expressing animals.

Prior to this work, an astrocyte-specific model of mtFUS expression had not been established, precluding evaluation of astrocyte-based non-cell autonomous effects *in vivo*. The novel model that we have developed demonstrates robust and reproduceable behavioral and pathological effects in a brief 2-week time frame. We anticipate that this method may be of considerable utility for others wishing to study astrocyte-mediated effects in neurodegenerative disease where mouse models are otherwise not available, due to rapid behavioral outcome measures and ease of testing therapeutic compounds. However, this system is not ideal for evaluating long-term effects or modeling aspects of FUS-ALS such as disease onset or progression, due to the very targeted and limited expression within the cervical spine region and adult age at which these animals underwent surgery. As such, while we are curious what the effects of a long-term continuous infusion of α-TNFα neutralizing antibody via an osmotic pump would be in the background of astrocytic mt-FUS expression, we feel that this investigation would be better suited for a chronic mtFUS expression mouse model, rather than the acute system that we have optimized.

Studies at the clinical, genetic, histological, and molecular levels support the hypothesis that neuroinflammation plays a key contributory role in ALS (McCauley & Baloh, 2019). Intriguingly, several previous studies investigating loss of functional FUS protein have suggested that FUS may have a significant role in regulating inflammatory signaling. B-cells derived from FUS-deficient mice have been shown to have reduced responses to traditional B-cell activating stimuli (Hicks et al., 2000). Additionally, FUS-deficient astrocytes display transcriptomic alterations that are enriched for inflammatory genes (Fujioka et al., 2013). Furthermore, while admittedly confined to a small sample size, reduction of FUS levels by 80% in the frontal cortex of marmosets for 6-8 weeks resulted in elevation of both GFAP and Iba1 positive cells and fluorescence intensity, resembling a state of reactive gliosis (Endo et al., 2018). In contrast, overabundance of FUS has also recently been implicated in sensitizing astrocytic and microglial responses to inflammatory environments. Cultured astrocytes overexpressing FUS *in vitro* display enhanced reactivity to inflammatory stimuli, and subsequently cause astrocyte-mediated motor neuron toxicity *in vitro* (Ajmone-Cat et al., 2019). While this complicated picture has emerged where either too little or too much FUS causes a reactive state in astrocytes, it is quite apparent that FUS levels and/or localization are tightly tied to signaling events indicative of cellular distress and inflammation.

At the cellular level, astrocytes act in a multi-faceted way to influence the inflammatory state of the CNS (Colombo & Farina, 2016). In the hippocampus, endogenous basal levels of astroglial-derived TNFα have been shown to be involved in synaptic scaling, whereby TNFα signaling leads to inclusion of calcium permeable GluR1 subunits within amino-3-hydroxy-5-methyl-4-isoxazolepropionic acid (AMPA) receptors in excitatory neurons, as well as decreasing the impact of inhibitory synaptic strength through selective endocytosis of aminobutyric acid type A (GABAA) receptors (Beattie et al., 2002, Santello et al., 2011, Stellwagen et al., 2005, Stellwagen & Malenka, 2006). Alternatively, using a mouse model of multiple sclerosis (experimental autoimmune encephalitis, EAE), where TNFα is elevated above homeostatic levels, it has also been shown that astrocytic TNFα signaling is responsible for NMDA receptor-mediated contextual learning/memory impairment (Habbas et al., 2015). Our previous studies investigating the effects of mtFUS astrocytes on wildtype endothelial cells of the blood brain barrier and wildtype spinal cord motor neurons both demonstrated that mutant FUS-expressing astrocytes display changes to genes involved in NF-κB and TNFα signaling pathways (Kia et al., 2018, Qosa et al., 2016), which are known to be potent regulators of inflammatory signaling (Kalliolias & Ivashkiv, 2016). In our *in vitro* co-culture model of mtFUS astrocytes and wildtype motor neurons, we have observed that with highly elevated TNFα levels, motor neuron AMPA receptor subunit composition is altered in a manner which renders these cells more susceptible to neuronal excitotoxicity (Kia et al., 2018). Further supporting this notion, work by Tolosa et. al. has shown that application of exogenous TNFα in the more complex system of organotypic rat spinal cord cultures also evoked glutamate-mediated excitotoxicity, through downregulation of astrocytic excitatory amino acid transporter 2 (EAAT2, also known as glutamate transporter 1 GLT1), elevation of extracellular glutamate levels, and activation of downstream NF-κB pathway oxidative stress responses (Tolosa et al., 2011). In our present work, we similarly demonstrate elevated TNFα mRNA and protein in the spinal cords of mice expressing mtFUS predominantly in astrocytes. We also observe changes to the astrocyte reactivity marker GFAP at the mRNA and protein levels by qRT-PCR and immunohistochemical staining. Additionally, our current work further supports this concept that astrocyte-secreted TNFα is a strong driving factor of motor neuron cell death in FUS-ALS, as expression of mtFUS of animals lacking TNFα did not display deficits in motor behavior or loss of cells in the spinal cord. Finally, our therapeutic intervention strategies targeting soluble TNFα in wildtype (TNF^+/+^) animals also suggest that lowering TNFα in FUS-ALS patients may be beneficial.

Evaluation of TNFα in ALS patients has consistently revealed elevated levels of both membrane bound and soluble TNFα and its receptors. This has been verified in plasma and cerebrospinal fluid (Cereda et al., 2008, Poloni et al., 2000), in post-mortem spinal cords by immunostaining (Kiaei et al., 2006), and in mRNA assessment from post-mortem spinal cord samples (Brambilla et al., 2016). A recent study using next generation RNA sequencing from cervical spinal cord of ALS patients also shows elevation of genes involved in inflammatory processes, with TNFα implicated as the main regulatory molecule for these genes (Brohawn et al., 2016). However, these studies which have examined primarily sporadic ALS patients, have shown that the extent of activation of the NF-κB and TNFα pathways have not correlated with disease duration or severity (Lu et al., 2016, Tateishi et al., 2010). Another layer of complexity also muddies interpretation of TNFα/NF-κB levels from patient samples. In addition to activating downstream signaling cascades, TNFα and NF-κB are known to participate in positive feedback loops where NF-κB can itself regulate TNFα transcription, and TNF signaling can act both in autocrine and paracrine manners to trigger NF-κB activation within an activating cell and in neighboring cells respectively (Gane et al., 2016, Pekalski et al., 2013). Overall, while some of the present evidence does suggest that elevated TNFα levels in ALS are likely a non-specific and self-perpetuating astrocytic and microglial response to degeneration of motor neurons, other studies have shown that there is a potential correlation between incremental increases in cytokine levels and disease progression (Tortarolo et al., 2017).

A different picture emerges when one focuses more specifically on the genetic forms of ALS. As was previously seen for sporadic cases, NF-κB levels are elevated in both astrocytes and microglia in the mouse model expressing pathogenic mutant SOD1^G93A^ mutation (Frakes et al., 2014). Further studies in these animals revealed that inhibiting NF-κB solely in astrocytes did not increase motor function or survival, nor did it alter disease progression or onset (Crosio et al., 2011). Preventing NF-kB signaling in microglia however did have a profound effect, preventing motor neuron cell death and extending survival through a mechanism involving reduced release of pro-inflammatory molecules (Frakes et al., 2014). Intriguingly, ablating the TNFα gene completely in SOD1^G93A^ or SOD1^G37R^ mice did not affect levels of spinal cord astrocytic or microglial reactivity, axon degeneration phenotypes, motor neuron loss, or overall animal survival (Gowing et al., 2006). Therefore, it appears that in the context of SOD1-ALS, that microglia are the major contributing glial population to NF-κB-mediated motor neuron toxicity, but that this effect is due to components of the NF-κB signaling cascade other than TNFα. It is likely therefore, that individuals with SOD1-ALS would not benefit as strongly to TNFα targeting therapeutics. It is not surprising that with different forms of genetic ALS, the mechanisms of non-cell autonomous toxicity may differ, given that different cellular effectors are perturbed as a result of altered protein localization and function. The NF-κB pathway has also been implicated in several models of TDP-43 based ALS, with enhanced NF-κB activation and upregulation of multiple downstream inflammatory cytokines including TNFα (Swarup et al., 2011). In contrast to overall NF-κB pathway upregulation, in both *in vitro* and *in vivo* models of mtFUS-ALS, we have now identified and verified TNFα-mediated motor neuron toxicity and suggest that targeting the TNFα may be a particularly promising avenue in genetic patients harboring these mutations (Kia et al., 2018, Qosa et al., 2016).

TNFα exerts its effects on cells through the actions of two ubiquitously expressed surface receptors, termed TNF receptor 1 (TNFR1) and TNF receptor 2 (TNFR2) with TNFR1 preferentially activated by soluble TNFa (sTNFα) and TNFR2 activated by membrane bound TNFα (mTNFα) respectively (McCoy & Tansey, 2008). Signaling through these receptors involves differing intracellular cascades and genes involved in either pro- or anti-apoptotic events, as well as pro- or anti-inflammatory signaling. TNFα can therefore alternatively promote protective or toxic effects depending on which receptor is activated. Two recent reviews on this topic are those of Guidotti (Guidotti et al., 2021) and Tortarolo (Tortarolo et al., 2017), which cover the differences in these pathways in much more detail as well as evidence for involvement of these specific receptors in SOD1-ALS mouse models. In our present *in vitro* and *in vivo* mtFUS models, we have not yet evaluated the contribution of specific TNFα receptor signaling in the neurotoxic effects that we have seen. Ongoing efforts to explicitly target sTNFα or mTNFα, and evaluating cells derived from TNFR receptor knockout animals will shed more light on the cellular mechanisms and downstream pathways involved.

Our results in TNFα knockout animals and effective blocking of motor neuron toxicity through use of an α-TNFα neutralizing antibody, hint at the potential utility of therapeutically targeting this signaling pathway in FUS-ALS. The testing of TNFα modulating therapeutics in ALS is not a novel concept, and several clinical trials have been run in the past. The first approach was using generic anti-inflammatory agents to inhibit TNFα synthesis. Promising results in SOD1^G93A^ mice showed that oral administration of the compound thalidomide improved motor behavior, increased motor neuron survival, delayed disease onset, and slowed disease progression (Kiaei et al., 2006). When TNFα immunoreactivity was examined in the spinal cord, the compound appeared to have the intended effect of destabilizing TNFα mRNA, thus lowering its expression. Unfortunately, when this went to a phase II clinical trial for safety and efficacy in humans, secreted TNFα levels were not impacted, the compound was not well-tolerated at the intended dosing, and disease progression was unaltered (Stommel et al., 2009). Similarly in a Phase III clinical trial, minocycline, which also inhibits TNFα synthesis and improved survival in pre-clinical mouse models, resulted in faster functional deterioration when tested in patients (Gordon et al., 2007). While not yet evaluated for use in ALS patients, an array of anti-TNFα molecules have been recently developed, tested, and approved for inflammatory disorders such as rheumatoid arthritis and Crohn’s disease, as well as for the neurodegenerative conditions multiple sclerosis and Alzheimer disease (Sedger & McDermott, 2014). These compounds, such as adalimumab and etanercept, do have serious adverse side-effects, which have been attributed to the non-specificity of action in inhibiting TNFα and general suppression of immune system function (Kemanetzoglou & Andreadou, 2017). An exciting new generation compound, XPro1595 selectively targets and neutralizes only sTNFα. In the EAE mouse model of multiple sclerosis, treatment with XPro1595 showed improved remyelination, preservation of axon integrity, and overall clinical outcome, while the non-selective TNFα inhibitor etanercept displayed no recovery of phenotype or function (Brambilla et al., 2011). Similarly, a mouse model of spinal cord injury has also shown that animals receiving XPro1595 demonstrated improved motor function and reduced spinal cord lesion damage, while animals receiving etanercept showed no improvement (Novrup et al., 2014). Importantly, these authors suggest that XPro1595 alters the inflammatory environment by neutralizing sTNFα, without suppressing neuroprotective effects of mTNFα signaling via the TNFR2 receptor (Novrup et al., 2014). Based on these preclinical findings, XPro1595 is eligible for testing in human diseases with known elevation of TNFα levels. In 2019, a Phase 1b trial was begun to assess safety and tolerability in patients with mild to moderate Alzheimer Disease (ClinicalTrials.gov Identifier: NCT03943264).

While further mechanistic investigation into the contributions of astrocytes to neuroinflammation in FUS-ALS are needed, our results do indicate that TNFα signaling is indeed perturbed *in vivo*. Additionally, our data strengthen the rationale for continued studies of cellular and molecular inflammatory mediators in sporadic ALS, as altered FUS localization or activity may also play a role in driving inflammation in ALS, even when mutated FUS is not the causative mechanism driving the disease (Tyzack et al., 2019). Finally, as medicine progresses towards an era of personalized therapy, it is optimistic to now envision tailored clinical trials to subsets of ALS patients. In our model we have identified TNFα as a promising therapeutic target for FUS-ALS patients. With newer generations of anti-TNFα compounds being generated to target soluble versus membrane TNFα as well as TNFR1 versus TNFR2 receptors, it is our hope that a targeted clinical trial for this subpopulation of genetic ALS patients would ultimately be warranted.

## MATERIALS AND METHODS

### Antibodies

Abcam: anti-FUS (Abcam Cat# 84078, RRID:AB_2105201, 1:1000), anti-GFP (Abcam Cat# ab13970, RRID:AB_300798, 1:1000); Cell Signaling: anti-NeuN (Cell Signaling Cat# 24307, RRID:AB_2651140, 1:400); Millipore: anti-ChAT (Millipore RRID:AB_2079751, 1:1,000),, anti-GFAP (Millipore Cat# AB5541, RRID:AB_177521, 1:500), anti-Olig2 (Millipore, RRID: AB_9610, 1:1000); Sigma: anti-MMP9 (Sigma-Aldrich Cat# M9570, RRID:AB_1079397, 1:60); Wako: anti-Iba1 (Wako Cat# 019-19741, RRID:AB_839504, 1:1000).

### Other Reagents

Agilent: Dako target antigen retrieval solution; AmericanBio: non-fat dry milk omniblock; Bio-Rad Laboratories: 4-20% Mini-PROTEAN TGX stain-free gel, broad-spectrum molecular weight ladder, Tris/glycine running buffer, Tris/glycine transfer buffer; Decon Laboratories: 200-proof ethanol; Electron Microscopy Sciences: 16% paraformaldehyde, Citifluor AF3; Millipore Sigma: β-mercaptoethanol, bovine serum albumin (BSA), Bradford reagent, deoxycholic acid, dithiothreitol (DTT), (ethylenedinitrilo) tetraacetic acid (EDTA), GeneRuler 1kb Plus, HEPES, hydrogen peroxide, immobilon-FL membranes, magnesium chloride hexahydrate (MgCl_2_), sodium dodecyl sulfate, sodium hydroxide; National Diagnostics: Histoclear; Research Products International: DEPC H_2_O; Thermofisher: agarose, Hoechst 33258 pentahydrate (bis-benzimide), methanol, normal goat serum, phosphate-buffered saline (PBS), potassium chloride, protease inhibitors, sodium chloride, Tris–HCl, Tween-20; US Biological: Nonidet (NP-40).

### Animals

Adult p180 female non-transgenic C57BL/6J mice were acquired from Jackson Laboratories (https://www.jax.org/strain/000664) and housed in a humidity-, temperature-, and light-controlled animal facility with *ad libitum* access to standard chow and water. For experiments involving TNFα knockout animals, the following genotype was also acquired from Jackson Laboratories: B6.129S-Tnf tm1Gkjl/J (https://www.jax.org/strain/003008). Homozygous female mice of this line were obtained at 12-weeks of age, which have a targeted deletion of the TNFα gene by replacement of 40 base pairs of the 5’ UTR, the ATG start codon, and the entire coding region for the first exon and part of the first intron with the MC1neopA cassette. 12-week-old female wildtype non-transgenic C57BL/6J mice as indicated above were acquired at the same time to serve as cohort controls, as the transgenic animals were also inbred using this background. Knockout and wildtype animals were aged to p180 in-house prior to intraspinal injection surgeries, to age-match timing with other surgical paradigms.

All experimental procedures were approved by the Thomas Jefferson University Institutional Animal Care and Use Committee (IACUC) and conducted in compliance with the National Institutes of Health Guide for the Care and Use of Laboratory Animals.

### Virus production

AAV9 vectors were generated by the University of Pennsylvania Penn vector core using a plasmid designed as follows. The gfa_104_ promoter element was from the pAAV.GFA104.PI.eGFP.WPRE.bgH plasmid, which was a gift from Philip Haydon (Addgene plasmid #100896). AAV9 gfa_104_-_eGFP::hFUS^R521G^ vectors were created by subcloning with a plasmid containing N terminally eGFP tagged human R521G (CGC to GGC) FUS in a pcDNA3.1/nV5 DEST backbone (Kia et al., 2018).

### Intraspinal delivery of AAV9 Virus

Intraspinal delivery of AAV9 in p180 mice was carried out as previously described (Li et al., 2015, Li et al., 2014). Briefly, deeply anesthetized mice underwent incision of their dorsal skin and underlying muscle, with retraction so that the spinous processes between C2 and T1 were visible. Bilateral laminectomy at spinal levels C3, C4, and C5 was then followed by six total injections, each containing a 1×10^11^ GC/injection of the AAV-gfa_104__eGFP viruses in 1 μl total volume using a Hamilton gas-tight syringe mounted on an electronic UMP3 micropump (World Precision International, Sarasota FL). A 33-gauge 45° beveled needle was lowered 0.8 mm below the dorsal surface of the spinal cord to target the ventral horns. Targeting of injections were guided along the lateral axis corresponding to the middle of each spinal segment, and along the rostral-caudal axis at the location of dorsal root entry for each bilateral injection set at levels C3, C4, and C5. Each injection was delivered over a 5-minute interval at a constant rate. Following completion of surgical procedures, overlying muscles were then closed in layers with sterile 4-0 silk sutures. The skin incision was closed using sutures followed by sterile wound clips. Animals recovered on a heating pad until awake and then were returned to their home cages. To minimize pain and discomfort, animals were administered subcutaneous sterile saline fluids, buprenorphine analgesic at 0.05 mg/kg, and cefazolin antibiotic at 10 mg/kg at the time of surgery, and at 12-hour intervals for 24 hours following surgery.

### Intraspinal delivery of anti-TNFα antibody

Our previous *in vitro* experiments demonstrated efficacy of targeting astrocytic-derived TNFα to prevent motor neuron toxicity using a TNFα neutralizing antibody (Kia et al., 2018). In the present study, functional grade anti-TNFα (α-TNFα, MP6-XT22, Thermofisher) antibody was used. In our first therapeutic paradigm, antibody was co-injected with AAV at the time of surgeries, at the previously demonstrated effective dosing of 2mg/kg, while maintaining the same 1uL total injection volume and viral titer. This antibody has been previously shown in rodent models to be safe, effective, and well-tolerated at this dosing (Finsterbusch et al., 2016, Via et al., 2001). Assessment of motor behavior and motor neuron numbers were carried out 2 weeks following surgery to determine potential beneficial effects of this therapeutic intervention.

In our second therapeutic paradigm, intraspinal delivery of AAV9 Virus was performed as indicated, with a non-adhering dressing (Adaptic non-adhering dressing by Systagenix) applied over the spinal cord prior to surgical closure. Following one week of viral expression, animals were again deeply anesthetized, their original surgical incision re-opened, and the spinal cord was again exposed by retraction of dorsal skin and underlying muscle. The non-adhering dressing was gently removed to re-expose the spinal cord. A pre-saturated gelfoam sponge (Gelfoam Dental sponges from Pharmacia & Upjohn Company) was directly applied to the exposed spinal cord region. Animals in this grouping either received gelfoam that had been saturated with functional grade anti-TNFα antibody at the 2mg/kg dosing, with sterile saline solution making up the remaining liquid volume, or a sterile saline saturated gelfoam as a control. Following completion of this second surgical procedure, overlying muscles were again closed with sterile 4-0 silk sutures. The skin incision was closed using sutures followed by sterile wound clips. Animals recovered on a heating pad until awake and then were returned to their home cages. To minimize pain and discomfort, animals were administered subcutaneous sterile saline fluids, buprenorphine analgesic at 0.05 mg/kg, and cefazolin antibiotic at 10 mg/kg at the time of surgery, and at 12-hour intervals for 24 hours following surgery. Assessment of motor behavior, and motor neuron numbers were carried out 2 weeks and 2 months following surgery to determine potential beneficial effects and longevity of this therapeutic intervention.

### Motor Behavioral Assessments

Wirehang latency to fall was measured by placing mice upside down on a metal wire grid at a height of 50 cm above a soft landing pad. The amount of time spent hanging onto the cage prior to falling was measured, with a maximum cutoff of 120 seconds. Muscle grip strength was determined for combined forelimbs using a DFIS-2 Series Digital Force Gauge (Columbus Instruments). Grip strength testing was performed by allowing the mice to grasp a thin “Y” bar attached to the force gauge. This was followed by pulling the animal away from the gauge until the forelimbs simultaneously released the bar. Force measurements were averaged from five trials for each animal.

### Spinal Cord Preparation for Immunofluorescence and qPCR

Mice were euthanized by carbon dioxide asphyxiation and whole animals were fixed using standard procedures. Briefly, a perfusion needle connected to a peristaltic pump was placed in the posterior end of the left ventricle of the heart. Mice were first perfused with approximately 200 mL phosphate buffered saline followed by 200 mL of 4% paraformaldehyde. Whole spinal cords were dissected and placed in 4% paraformaldehyde at 4 °C for 30 minutes, then moved to 2% paraformaldehyde overnight. Paraformaldehyde was rinsed off the tissue with 3 consecutive washes with a 1:1 solution of potassium monosulfate and potassium bisulfate buffer. Spinal cords were then placed in 30% sucrose for 24-48 hours until tissue sinking. Following anatomical assessment, cervical spinal cord regions were frozen into Tissue-Tek OCT solution and sectioned at 30 μm using a cryostat (Thermo Scientific) and placed onto charged glass slides.

For tissue immunofluorescence, blocking, permeabilization, and staining conditions were carried out according to standard protocols and based on the manufacturer instructions for each antibody. Slides containing 30 µm cervical spinal cord tissue sections were heated overnight at 55°C, followed by Histoclear and rehydration in sequential 100, 95, 90, and 70% ethanol washes. Incubation in hydrogen peroxide (3% in methanol) for 30 minutes was used to block endogenous peroxidase activity. Unmasking of antigens was achieved by incubation in target antigen retrieval citrate solution for 1 hour at 95°C. Sections were next blocked in normal goat serum (10% in PBS) for 1 hour at room temperature (5% BSA instead of goat serum for ChAT) and incubated in primary antibodies overnight at 4°C (48 hour incubation specifically for ChAT). Primary antibodies included: anti-ChAT (Millipore RRID:AB_2079751, 1:1,000), anti-FUS (Abcam Cat# 84078, RRID:AB_2105201, 1:1000), anti-GFAP (Millipore Cat# AB5541, RRID:AB_177521, 1:500), anti-GFP (Abcam Cat# ab13970, RRID:AB_300798, 1:1000), anti-MMP9 (Sigma-Aldrich Cat# M9570, RRID:AB_1079397, 1:60), anti-NeuN (Cell Signaling Cat# 24307, RRID:AB_2651140, 1:400), anti-Iba1 (Wako Cat# 019-19741, RRID:AB_839504, 1:1000), and anti-Olig2 (Millipore, RRID: AB_9610, 1:1000). Following PBS washing, secondary labeling with AlexaFluor488, AlexaFluor594, or AlexaFluor647 (Life Technologies) were used to enable visualization. Hoechst stain (ThermoFisher) was used to label nuclei. Stained slides were mounted in Citifluor AF3 (Electron Microscopy Sciences). Image capture and processing were accomplished using a Nikon A1+ confocal microscope with NIS-Elements software. ChAT^+^ and MMP^+^ cells were assessed using bilateral ventral horn images with manual counting.

For qPCR analysis, RNA was isolated from fixed spinal cord sections using the PureLink RNA Mini Kit (ThermoFisher), reverse transcribed using the QuantiTect Reverse Transcription Kit (Qiagen), prepared for qPCR using a SYBR Green qPCR master mix (ThermoFisher) and analyzed on QuantStudio 5 Real-Time PCR System (ThermoFisher), Samples were measured in triplicate from each animal for each gene of interest. Data were normalized using GAPDH levels, fold change between groups is represented. Primer pairs obtained from Integrated DNA Technologies (Coralville, IA) for each gene are listed in Table 1.

**Table 1:**
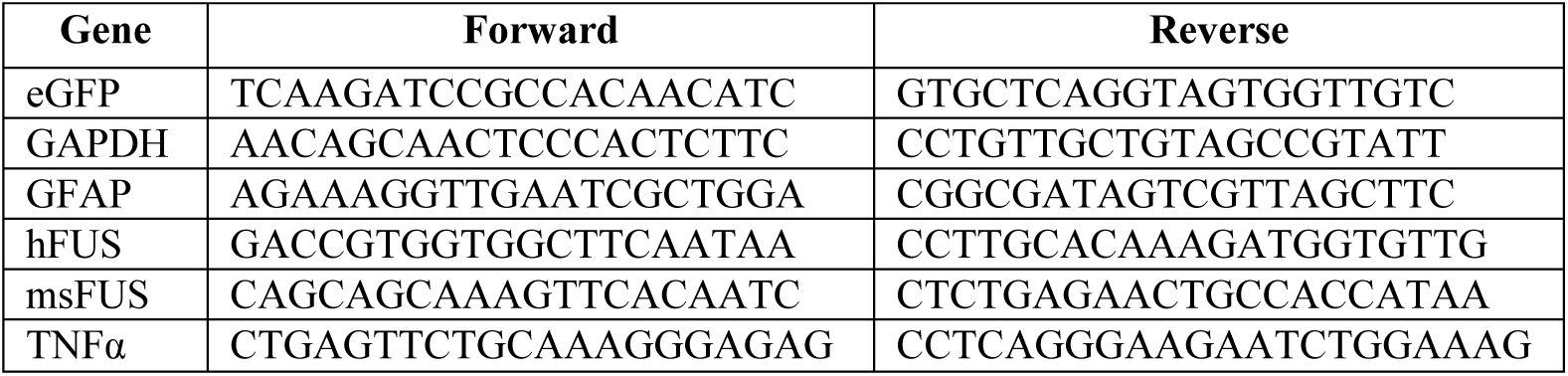
Primer pairs used for qPCR analysis of mouse spinal cord tissue samples

### Spinal Cord Preparation for ELISA and Western Blotting

Mice were euthanized by carbon dioxide asphyxiation. Cervical spinal cords were then dissected, and flash frozen in liquid nitrogen. Dissociated single-cell suspensions were prepared using homogenization buffer containing: 10 mM HEPES pH 7.9, 1.5 mM MgCl2, 10 mM KCl, 0.5 mM DTT, and protease inhibitor cocktail (1:100), and then underwent centrifugation at 845 g at 4°C for 5 minutes.

### TNFα ELISA

The supernatant from this homogenization step was used to evaluate levels of TNFα using the mouse TNF alpha High Sensitivity ELISA Kit according to manufacturer’s instructions (Invitrogen, BMS607HS). Briefly, microwell strips pre-coated with anti-TNFα capture antibody were washed with Wash Buffer. Samples and standards were added to wells along with Biotin-Conjugate and allowed to incubate for 2 hr. Wells were washed and then incubated with Streptavidin-HRP for 1 hr. Following another round of washing, wells were incubated next with Amplification Solution I for 15 min, were washed again, and then incubated with Amplification Solution II for 30 min. After a final washing step, wells were incubated with a TMB substrate solution for 15 min. All incubations were performed with shaking at room temperature. The Stop Solution was added immediately prior to plate reading. The color intensity of each well was measured at 450 nm using a Cytation5 microplate reader (Biotek).

### Western Blotting

From each animal, dissociated cells were then resuspended in RIPA buffer containing: 150 mM NaCl, 1% Nonidet (NP-40), 0.5% deoxycholic acid, 0.1% sodium dodecyl sulfate, 50 mM Tris–HCl, 2 mM EDTA, and protease inhibitor cocktail (1:100)). These suspensions were pulse-sonicated 3 times for 10 seconds each with 2 minutes rest on ice between rounds, and spun at 21,130 g at 4°C for 30 minutes. The Bradford method was used to determine protein concentrations of supernatant fractions. 25 μg of protein was loaded onto a Mini-PROTEAN TGX stain-free gel (4-20%) for electrophoretic separation, along with a molecular weight ladder. Following separation, total protein levels were determined by imaging with the pre-set stain-free gel parameters on the Bio-Rad ChemiDoc Touch Imaging System. Proteins were transferred onto Millipore Immobilon-FL membranes and were blocked in PBST (PBS + 0.1% Tween-20) and milk (5%) for 30 minutes at room temperature prior to overnight incubation at 4°C with primary antibodies in PBST + 5% milk. Primary antibodies used were: anti-FUS (Abcam Cat# 84078, RRID:AB_2105201, 1:1000)), and anti-GFP (Abcam Cat# ab13970, RRID:AB_300798, 1:1000. Three PBST washes were followed by incubation with LiCor antigen species-specific fluorescent probe-conjugated secondary antibodies (1:15,000 dilution in PBST + 5% milk) for 1 hour at room temperature. An Odyssey Infrared Imaging system (LiCor) equipped with ImageStudio was used for membrane visualization. Bands of interest were selected at the heights corresponding to the molecular weight of the protein of interest using ImageJ software. Pixel intensities of these defined areas were normalized to a horizontal section of the Bio-Rad total protein image spanning a broad molecular weight range. As such, overall intensity of each lane was determined to account for variability in tissue lysate extracts.

### PCR Analysis of TNFα knockout animals

Ear snips were collected at the time of mouse euthanasia for the TNF KO and wildtype paired experiments. DNA was isolated from these samples as follows. 75 μL of 1x Solution A was added to tissues, which were then heated to 95 °C for 30 minutes and then cooled to 4 °C. After mixing, 75 μL of 1x Solution B was added and mixed. Ingredients for solutions found in Table 2.

**Table 2:**
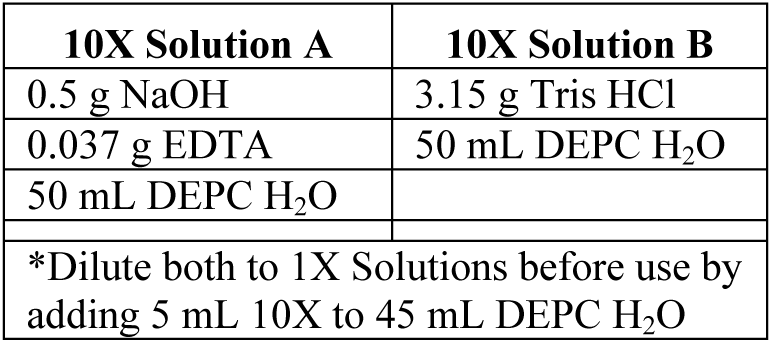
Solutions used to extract DNA for PCR analysis

Genotyping was performed using the suggested primer pairs provided by the Jackson Laboratory for these animals (below), along with the associated recommended PCR cycling times and temperatures (https://www.jax.org/Protocol?stockNumber=003008&protocolID=22433). PCR results were run on a 1.5% agarose gel, along with a DNA ladder (GeneRuler 1kb plus). Primer pairs obtained from Integrated DNA Technologies (Coralville, IA) are listed in Table 3.

**Table 3:**
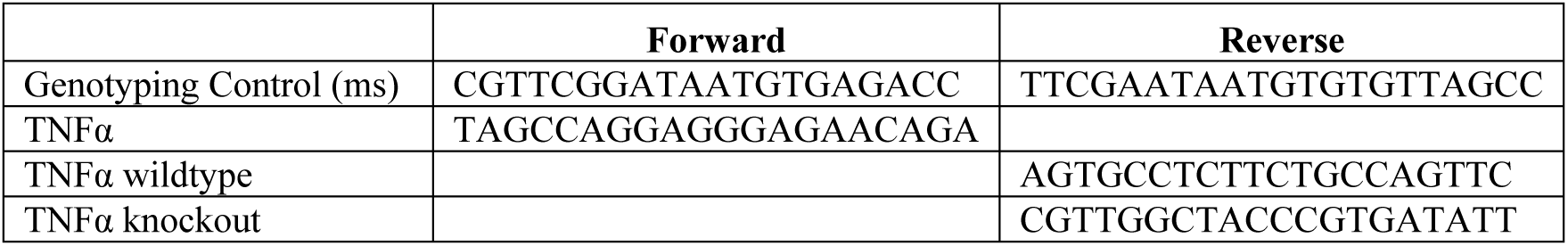
Primer pairs used for PCR analysis of wildtype and TNFα knockout animals

### Statistical Analysis

Prism 8.0 Software (GraphPad) was used to perform all statistical analyses. Statistical significance for pairwise assessment in experiments with multiple groups was accomplished using One-way ANOVA with post hoc Dunnett’s multiple comparison test or Sidak’s multiple comparison test where applicable. Student’s *t* test was used to evaluate any direct comparison of only two groups. All data are expressed as mean ± SEM, with values of *p* < 0.05 considered significant.

### Data Availability

No large scale data sets have been generated in this study.

## AUTHOR CONTRIBUTION

Conceptualization: B.K.J.; K.M.; P.P;

Methodology: B.K.J.; K.M.; N.M.H.; A.C.L;

Investigation: B.K.J; K.M.; N.M.H;

Formal Analysis: B.K.J; K.M.;

Visualization: B.K.J; K.M.; P.P.; D.T.; A.H.; H.I.;

Writing – Original Draft: B.K.J.; P.P;

Funding Acquisition: P.P.; D.T.; A.H.; A.C.L.;

Resources: P.P.; D.T.; A.H.; H.I.; A.C.L.;

Supervision: P.P.; D.T.; A.H.; H.I.

## THE PAPER EXPLAINED

### Problem

Mutations in DNA/RNA binding protein Fused in Sarcoma (FUS) have been implicated in approximately 4% of familial and 1% of sporadic cases of amyotrophic lateral sclerosis (ALS). Normally this protein is involved in many cellular functions due to binding with target DNA and RNA, including essential cellular processes of transcription, RNA processing, translation, and DNA damage repair. However, ALS-causing mutations frequently result in cytoplasmic mislocalization and aggregation of FUS protein, disrupting these critical processes. These aggregates occur not only in the motor neurons of the spinal cord which are cells that die in ALS, but also in the surrounding glial support cells. Growing evidence has suggested that glial cells also are important drivers of disease in other genetic forms of ALS. While studies have investigated the effects of FUS mutations within neuronal cells, much less is currently known about potential deleterious consequences to motor neurons arising from neighboring astrocytes expressing mutant FUS protein.

### Results

Here, we have developed an *in vivo* model for expressing mutant FUS protein in the cervical spinal cord of adult mice. Following two weeks of such expression, animals display deficits in motor function by behavioral tests, and have reduced motor neuron numbers within their spinal cords. In agreement with our previous *in vitro* studies, we validated that the pro-inflammatory cytokine TNFα was elevated following mutant FUS expression in astrocytes. We next confirmed that a major component of the observed *in vivo* toxicity was TNFα−dependent, as TNFα knockout animals did not have any motor phenotypes or neuronal cell loss. Finally, when we used an anti-TNFα-neutralizing antibody to prevent TNFα-mediated signaling from occurring, we were successful at preventing astrocyte-derived toxicity. Importantly, this beneficial effect was seen either if the neutralizing antibody was given at the time when mutant FUS was just beginning to be expressed, or if we waited and allowed mutant FUS to have cellular effects for one week prior to therapeutic intervention.

### Impact

Identifying and targeting astrocytic contributions driving ALS pathogenesis is a potential strategy for therapeutic intervention in patients. We have verifed TNFα as a *bona fide* contributor to motor neuron loss in FUS-ALS and have demonstrated that targeting this molecule is possible therapeutically. In doing so we have set the stage for further investigation of TNFα pathway modulators as potential therapeutic strategies for FUS-ALS, and perhaps more broadly to sporadic ALS cases where mutant FUS is also frequently found in cytoplasmic inclusions.

## FOR MORE INFORMATION

https://www.jefferson.edu/university/farber_institute/weinberg_als_center.html

http://www.alscenter.org/

http://www.alsa.org/

**EV1: Additional characterization of mouse model**

A) Illustration of how masking was performed to enable quantification of GFP^+^ cells in spinal cord sections. These images show a mask of thresholded DAPI signal (left) and a mask of thresholded eGFP:hFUS^R521G^ signal (right), performed on 20x images.

B) Localization pattern of eGFP:hFUS^R521G^ (mtFUS), showing a strong nuclear expression pattern along with cytoplasmic aggregates, 60x magnification, scale bar indicates 20 μm.

C) Animals were monitored following viral injections to assess if any overt weight loss was observed. At four weeks post-injection, no differences were noted in the eGFP versus mtFUS cohorts. n = 4 mice per group were evaluated. Data presented as mean ± SEM.

**EV2: eGFP and FUS expression levels are comparable in GFP and mtFUS animals**

A) Immunohistochemical staining was performed to evaluate GFP expression levels with the ventral horn of eGFP and mtFUS expressing animals at 3-days (top) and 14-days (bottom) post injection. Representative 20x images reveal similar numbers of transfected cells and overall GFP levels. Scale bar indicates 250 μm.

B) qPCR analysis targeted for evaluating eGFP levels at 14 days post-injection indicates a robust and comparable level of expression in GFP and mtFUS expressing animals compared with the sham surgery group. Data presented as mean ± SEM, Dunnett’s multiple comparison test determined statistical significance, n=4 animals per group, ***p< 0.001.

C) Lysates from fresh frozen C4-C6 spinal cord lysates from eGFP and mtFUS animals were generated. These lysates were immunoblotted for GFP protein and normalized to total protein loading. Quantification of band intensities revealed equivalent levels of GFP protein expression. Only the mtFUS expressing animals showed a GFP positive band at the anticipated 100kD height of the eGFP:hFUS^R521G^ fusion protein. Data presented as mean ± SEM, n=6 animals per group, student’s t test determined statistical significance.

D) Immunohistochemical staining was performed to evaluate total FUS expression levels with the ventral horn of eGFP and mtFUS expressing animals at 14-days post injection. Representative 20x images reveal similar overall FUS levels (shown in red), scale bar indicates 250 μm.

E) qPCR analysis targeted for evaluating mouse (left) and human (right) FUS levels at 14 days post-injection indicates a slight but not significant reduction in mtFUS expressing animals. However, a specific, significant level of human FUS expression is only seen in the mtFUS expressing animals. Data presented as mean ± SEM, Dunnett’s multiple comparison test determined statistical significance, n=4 animals per group, **p< 0.01.

F) Western blot analysis of fresh frozen C4-C6 spinal cord lysates from eGFP versus mtFUS animals revealed equivalent levels of FUS protein expression. Again only the mtFUS expressing animals showed a higher weight band at the anticipated 100kD height of the eGFP:hFUS^R521G^ fusion protein. Data presented as mean ± SEM, n=6 animals per group, student’s t test determined statistical significance.

**EV3: Motor neuron numbers are not altered 3-days post virus injection**

A) Left: Representative 60x magnification immunohistochemical staining for MMP9 in spinal cord sections from eGFP and mtFUS expressing animals at 3 days post-injection, scale bar indicates 125 μm. eGFP and mtFUS expressing animals show similar numbers of MMP9^+^ neurons at this timepoint, visualized in cyan. Right: quantification of MMP9^+^ motor neurons was performed as in Figure 2. Analysis revealed comparable numbers of MMP9^+^ cells in sham surgery, eGFP and mtFUS cohorts. Data presented as mean ± SEM, n = 3 animals per condition, m= 4 ventral horns per animal. Sidak’s multiple comparison test determined statistical significance.

B) Left: Representative 60x magnification immunohistochemical staining for ChAT at 3 days post-injection in eGFP and mtFUS expressing animals, scale bar indicates 125 μm. At this timepoint, eGFP and mtFUS expressing animals show similar numbers of ChAT^+^ neurons visualized in magenta and indicated by *. Right: quantification of ChAT^+^ motor neurons was performed as in Figure 2, and similar numbers of ChAT^+^ cells were found across the three experimental groups. Data presented as mean ± SEM, n = 3 animals per condition, m= 4 ventral horns per animal. Sidak’s multiple comparison test determined statistical significance.

**EV4: mtFUS animals display elevated astrocyte reactivity**

A) qPCR analysis targeted for evaluating glial fibrillary acidic protein (GFAP) levels at 14 days post-injection indicates a significantly elevated level of mRNA expression in mtFUS expressing animals compared with sham surgery and eGFP control groups. Data shown as relative fold-changes compared to controls, and are presented as mean ± SEM, Student’s t test determined statistical significance, n=5 animals per group, \*\*\**p* < 0.01.

B) Representative 20x magnification immunohistochemical staining for GFAP, co-labelled with GFP to visualize transduced cells at 14 days post-injection, scale bar indicates 125 μm. mtFUS expressing animals show elevated GFAP fluorescence intensity compared to eGFP expressing animals.

**EV5: TNFα wildtype and knockout animals have been validated by PCR analysis and express mtFUS to comparable levels**

A) PCR analysis revealed correct genotype band patterns for wildtype (WT) and TNFα knockout (KO) animals, based on suggested primers obtained from the Jackson laboratory. A common forward primer was used, with specific reverse primers for either the wildtype or mutant DNA sequences. The mutant allele amplifies at a size of 318 base pairs, and the wildtype amplifies at 183 base pairs. A control PCR for mouse DNA was also included with an amplification size of approximately 400 base pairs. All knockout animals were homozygous for the mutant allele, showing only a single band at the 318 base pair height.

B) qPCR analysis targeted for evaluating eGFP levels at 14 days post-injection indicates a robust and comparable level of expression in wildtype and TNFα knockout animals expressing mtFUS compared with their respective sham surgery groups. Data presented as mean ± SEM, Sidak’s multiple comparison test determined statistical significance, n=6 animals per group, ***p< 0.001.

**EV6: mtFUS expression is comparable in animals receiving α−TNFα neutralizing antibody and animals receiving mtFUS alone**

A) In the paradigm of co-injection of α-TNFα neutralizing antibody at the time of surgery, qPCR analysis targeted for evaluating eGFP levels at 14 days post-injection indicates a robust and comparable level of expression in both mtFUS and mtFUS + α-TNFα animals. Data presented as mean ± SEM, One-way ANOVA with Sidak’s multiple comparison test determined statistical significance, n=8 animals per group, ** *p* < 0.001.

B) In the paradigm of viral infection with mtFUS, followed a week later by α-TNFα neutralizing antibody application, qPCR analysis was performed to evaluate eGFP levels at 14 days and 2 months post-injection. At both timepoints, a robust and comparable level of expression is seen in both mtFUS and mtFUS + α-TNFα animals. Data presented as mean ± SEM, One-way ANOVA with Sidak’s multiple comparison test determined statistical significance, n=4 animals per group, * *p* < 0.05.

## ACKNOWLEDGEMENTS

We would like to acknowledge members of the Jefferson Weinberg ALS Center for suggestions and critical evaluation of this work. We thank laboratory members Kristian Mayer and Thomas Westergard for assistance in preliminary testing of model development and behavioral endpoint testing. N-terminally eGFP tagged human FUS in the pcDNA3.1/nV5-DEST backbone was originally received as a kind gift from Dr. Aaron Gitler. This work was supported by funding from the NIH (R01 NS051488 and R01 NS109150 to P.P, R01 NS079702 and R01 NS110385 to A.C.L., RF1-AG057882 and R21-NS103118 to D.T., R00 NS091486 and R21 NS116761 to A.R.H.), the Muscular Dystrophy Association (D.T.), the Robert Packard Center for ALS Research (D.T.), the Family Strong 4 ALS foundation (B.K.J. and P.P), and the Farber Family Foundation (B.K.J. and P.P).

